# Site-, strain- and mutation-specific osteopetrosis phenotypes in mice with *Clcn7* variants

**DOI:** 10.64898/2026.10.31.763333

**Authors:** Anika Shimonty, Xenia G. Dahl, Elissa Zboinski, Dyann Segvich, Kelly Tsang, Emily McDonald, Joseph M. Wallace, Thomas L. Andersen, Julia F. Charles

## Abstract

**Background:** Autosomal dominant osteopetrosis type II (ADOII) is a rare bone disorder caused primarily by heterozygous mutations in CLCN7 and characterized by increased bone mass yet fragility and marked phenotypic heterogeneity among affected individuals. The biological factors underlying this variability remain poorly understood. We investigated the effects of age, sex, genetic background, and mutation type on bone phenotypes in murine models of ADOII carrying heterozygous *Clcn7* F318L/+ or G213R/+ mutations.

**Methods:** Bone microarchitecture was assessed by micro-computed tomography in vertebrae and femora from female and male mice across multiple ages (12, 24, and 52 weeks) and strain backgrounds (FVB/NJ, C57BL/6, and 129). Vertebral biomechanical properties were estimated by micro-finite element analysis, while femoral mechanical properties were measured using three-point bending.

**Results:** *Clcn7* F318L/+ mice exhibited significantly increased trabecular bone compared with wild-type littermates at vertebral and femoral sites across the lifespan in both sexes. The magnitude of the vertebral trabecular phenotype increased with age, whereas age-related effects in the femur were more modest and site dependent. Vertebral micro-finite element analysis predicted greater stiffness and ultimate force in F318L/+ mice irrespective of age. Cortical thickness was modestly increased, particularly in female mice, but femoral mechanical properties were largely unchanged. The relative increase in vertebral trabecular bone was similar on the FVB/NJ and C57BL/6 backgrounds in males but greater on C57BL/6 in females, whereas the femoral trabecular and cortical responses were markedly attenuated on C57BL/6 in both sexes. In G213R/+ mice, the relative trabecular phenotype was similar on the C57BL/6 and 129 backgrounds, except for a greater vertebral phenotype in 129 males. Likewise, comparison of F318L/+ and G213R/+ mice demonstrated comparable vertebral phenotypes, although mutation-specific differences emerged at the femur, particularly in males.

**Conclusion:** Collectively, these findings demonstrate that heterozygous *Clcn7* mutations produce a robust osteopetrotic phenotype characterized primarily by increased trabecular bone mass that is present regardless of sex, age, genetic background, and mutation type. Age and genetic background modify its magnitude in a site- and sex-specific manner, suggesting that local factors may contribute more strongly to phenotypic variation than the specific disease-causing *Clcn7* mutation. These results provide insight into determinants of phenotypic heterogeneity in ADOII and establish a framework for future mechanistic studies of osteopetrotic bone quality and fragility.

## Introduction

Osteopetrosis is a group of rare genetic bone disorders characterized by high bone mass yet paradoxically structurally brittle bones prone to fracture [1, 2]. This high bone mass arises from impaired osteoclastic resorption, caused either by defective osteoclast differentiation or dysfunction in osteoclast activity, while the bone formation is maintained. Based on inheritance patterns, three forms of osteopetrosis are recognized: Autosomal recessive osteopetrosis (ARO), Autosomal dominant osteopetrosis (ADO), and X-linked osteopetrosis [3].

Autosomal dominant osteopetrosis type II (ADOII), also known as Albers-Schonberg disease, is the most common form of osteopetrosis, with an estimated incidence of 1 in 20,000 [1]. ADOII primarily results from heterozygous mutations in the chloride channel 7 (CLCN7) gene [4–6]; however, there have been rare cases with T-cell immune regulator 1 (*TCIRG1*) mutations [7, 8]. The major clinical characteristics of ADOII patients include osteosclerosis and the distinctive radiographic ‘bone-in-bone’ appearance [9]. The presentation of ADOII is highly variable and may include increased bone fragility leading to fractures, including recurrent fractures, impingement of cranial nerves, encroachment of osteopetrotic bone into the marrow space, poor bone vascularity, and osteomyelitis [9–11]. Notably, disease severity and onset differ widely, even among patients from the same family and carrying the same mutation [4, 12]. At present, there is no disease-specific treatment for ADOII.

*CLCN7* encodes a lysosomal Cl−/H+ antiporter that functions as a dimer to mediate the exchange of chloride ions and protons [13]. *CLCN7* is expressed ubiquitously in different tissues [14, 15]. In osteoclasts, CLCN7 is localized to the ruffled border, where it is necessary for the acidification of the resorption lacuna and subsequent bone resorption [16]. Mutations in *CLCN7* impair chloride transport required for lacunar acidification, leading to osteoclast-rich osteopetrosis characterized by normal osteoclast numbers but defective resorptive function [2].

More than thirty-four different mutations in *CLCN7* have been reported in ADOII patients [17]. Among them, a p.G215R mutation was discovered in multiple patients [5], and later found to be one of the most frequent [18]. The orthologous p.G213R mutation was introduced into the mouse genome, generating the first murine model of ADOII [18]. Heterozygous p.G213R mutation results in a heterodimer protein that is retained in the endoplasmic reticulum due to a trafficking defect [19]. Another less common *CLCN7* mutation in ADOII is p.F318L, and a mouse model with the orthologous p.F316L mutation has also been generated; by convention, this mouse model is referred to as F318L [20]. In contrast to the p.G213R mutation, the heterodimeric protein resulting from the p.F316L mutation has no trafficking defect, but a functional defect resulting in inhibition of ion transport [20]. In both mouse models, homozygous mutation results in a severe form of disease and death early in life, while heterozygous mutation results in an ADOII-like phenotype.

Bone phenotypes are determined by multiple biological and environmental factors. Genetic factors are a major determinant of bone phenotypes, although they are not the only determinants. Genetic background, sex, age, diet, physical activity, and environment all modulate bone health [21–25]. Age profoundly influences skeletal biology [26–28]. Bone mass accrues during growth, but trabecular bone begins to decline before midlife, followed later by cortical bone loss, contributing to age-related fragility [26–28]. Aging is characterized by increased endocortical resorption and cortical porosity along with decreased trabecular and cortical bone mineral density collectively increasing fracture risk [28]. Sex also modifies skeletal structure. The skeleton is sexually dimorphic in both humans and mice, with females generally having more rapid bone loss than males [29, 30], accompanied by distinct gene expression profiles [30–32]. In mice, genetic background strongly influences bone microarchitecture and strength, with strain-dependent baseline differences, similar to the genetic diversity in humans [33–36] which may contribute to the marked phenotypic heterogeneity observed in ADOII patients.

In our present study, we investigated the impact of age, sex, genetic background, and specific *Clcn7* mutations on the bone phenotype in mouse models of ADOII. We used female and male mice of different ages, strain background, and two distinct *Clcn7* mutations to investigate the impact of these variables on bone phenotype and test the hypothesis that they would contribute to phenotypic heterogeneity. We performed micro-CT analysis of vertebra and long bones, as well as mechanical tests, to determine bone structure and biomechanical properties. Contrary to our hypothesis, the vertebral trabecular phenotype was robust. F318L/+ and G213R/+ mice of both sexes had increased vertebral BV/TV at every age and on every background examined. The relative increase did not differ between the two mutations, and genetic background influenced its magnitude. The femoral phenotype was more variable. Femoral trabecular bone was less influenced by age than vertebral bone, the femoral trabecular phenotype of F318L/+ mice was markedly attenuated on the B6 background in both sexes, and the increase was greater in G213R/+ than in F318L/+ males. Increased femoral cortical thickness was most evident in females and depended on genetic background. Mechanical properties were altered only in females, with higher yield force at 12 and 24 weeks of age and a stiffer but less ductile femur at 24 weeks.

## Results

### Increased trabecular bone in *Clcn7 F318L/+* mice across the age span

To understand the effects of aging on the *Clcn7* phenotype, we analyzed the 4th lumbar vertebra from 12-, 24-, and 52-week-old female wildtype (+/+) and F318L/+ littermate mice on the FVB/NJ background. There was no significant difference in either body weight or body length between the +/+ and F318L/+ mice at any age (Supplementary Table S1). Vertebral trabecular bone volume fraction (BV/TV) was higher in F318L/+ female mice at every age (Fig 1A, B). Consistent with this, F318L/+ mice had higher trabecular number (Tb.N), trabecular thickness (Tb.th), lower trabecular spacing (Tb.Sp), and higher bone mineral density (Tb.BMD) (Supplementary Table S2). Furthermore, finite element analysis (FEA) of vertebra shows that F318L/+ mice bones had higher predicted ultimate force and stiffness than their littermate +/+ controls, regardless of age (Supplementary Table S3).

**Figure 1:**
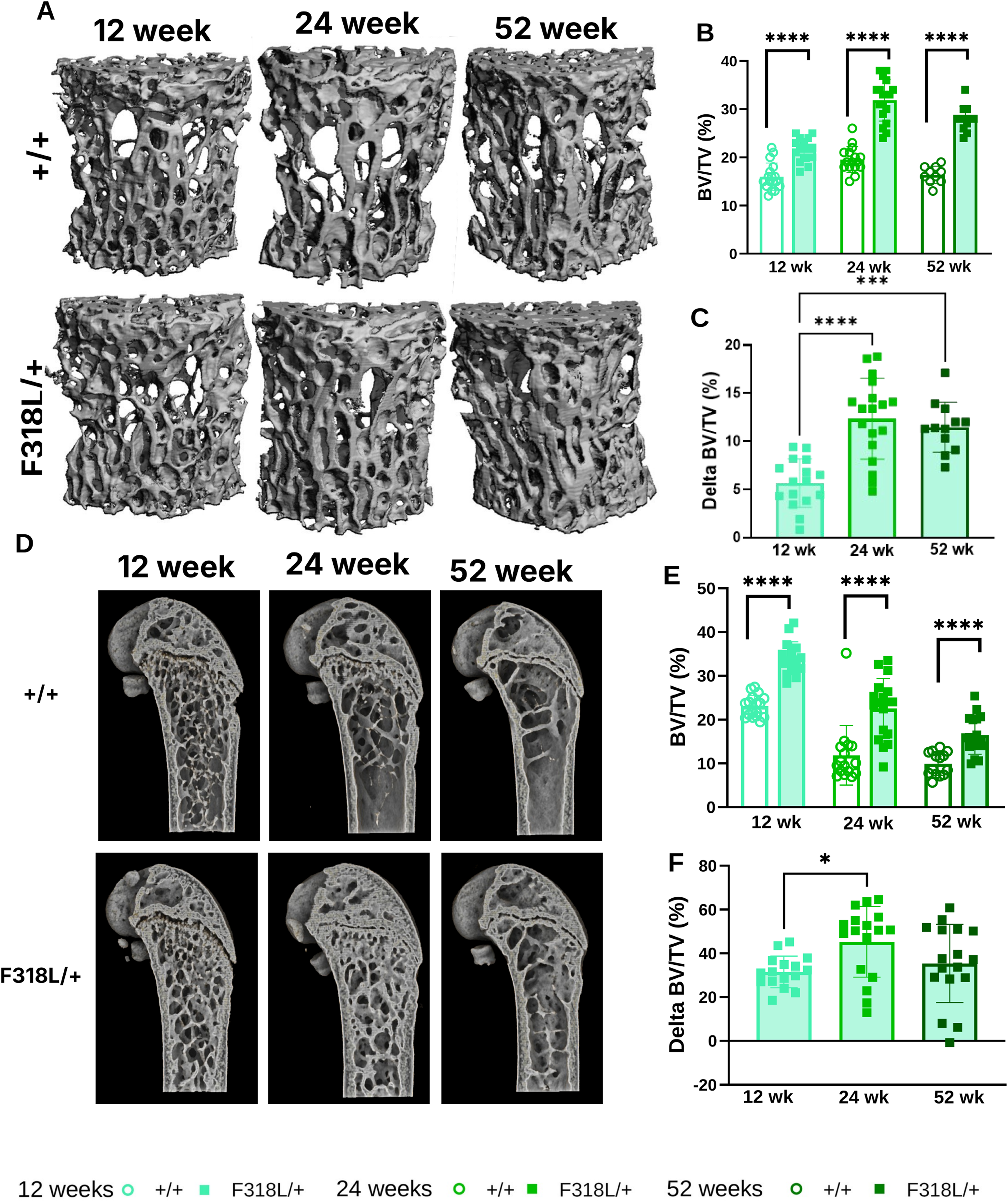
Vertebral trabecular BV/TV is increased in female F318L mouse model of ADOII. **A.** Representative 3D images of 12-, 24-, and 52-week-old female +/+ and F318L/+ vertebrae. **B.** Trabecular BV/TV from +/+ (open columns) and F318L/+ (filled columns) mice at 12-week (light green), 24-week (dark green), 52-week (Kelly green) of age. **C.** Δ BV/TV at each age. **D.** Representative 3D images of 12-, 24-, and 52-week-old female +/+ and F318L/+ femurs. **E.** Trabecular BV/TV, **F.** Δ BV/TV using the same color scheme as B-C. Data are presented as individual animals with mean ± SD. Δ (%) = (value of each mutant mouse − mean of the +/+ corresponding littermates) / mean of those +/+ littermates × 100. n= 15/group. +/+ vs F318L/+ at each age: Mann-Whitney test; Δ across ages: Kruskal–Wallis test with Dunn’s multiple-comparisons test. *= p<0.05, **= p<0.01, ***= p<0.001, ****= p<0.0001.

To determine if the severity of the phenotype varied with age, we calculated the Δ (delta) BV/TV of F318L/+ mice relative to the mean of their +/+ counterparts and compared ΔBV/TV across the age span. The impact of F318L/+ on vertebral BV/TV increased with age, as demonstrated by a greater Δ BV/TV in the 24- and 52-week-old compared to the 12-week-old cohort, although ΔBV/TV did not differ significantly between the 24- and 52-week-old cohorts (Fig 1C).

We next analyzed the trabecular compartment of femurs as a representative long bone. At all ages examined, female F318L/+ mice had significantly higher trabecular BV/TV (Fig 1D, E), with corresponding changes in Tb.N, Tb.Th, and Tb.S (Supplemental Table S4). Similar to vertebral trabecular bone, the Δ BV/TV of F318L/+ mice over their respective +/+ littermates increased with age between 12 and 24 weeks, though no further increase was observed at 52 weeks (Fig 1F), because femoral trabecular BV/TV declined significantly between 12 and 24 weeks of age in both +/+ and F318L/+ mice (Supplementary Table S4). This age-related loss of femoral trabecular bone contrasts with the vertebrae, where the mutant phenotype became more pronounced with age (Fig 1C).

To assess whether the impact of *Clcn7* F318L/+ on bone phenotype with aging is similar in both sexes, we analyzed male cohorts at 12-, 24-, and 52-weeks of age. Similar to females, the F318L/+ mutation increased trabecular BV/TV in males at all ages at both vertebrae (Fig 2A, B) and femur (Fig 2D, E) with no impact on body weight or length (Supplementary Table S1). As in females, the impact of F318L/+ on trabecular BV/TV measured as Δ BV/TV increased with age at the vertebrae, though this effect was less apparent at the femoral site (Fig 2C, F), where trabecular BV/TV declined with age in both genotypes (Supplementary Table S4). Overall, the impact of the *Clcn7* F318L mutation on trabecular bone was generally similar at both sites independent of sex. While age impacted the severity of the trabecular phenotype in both sexes at the vertebrae, age had a more modest impact on femoral trabecular bone in females and no impact in males, suggesting a site-specific effect of aging on the trabecular bone phenotype of *Clcn7* F318L mice.

**Figure 2:**
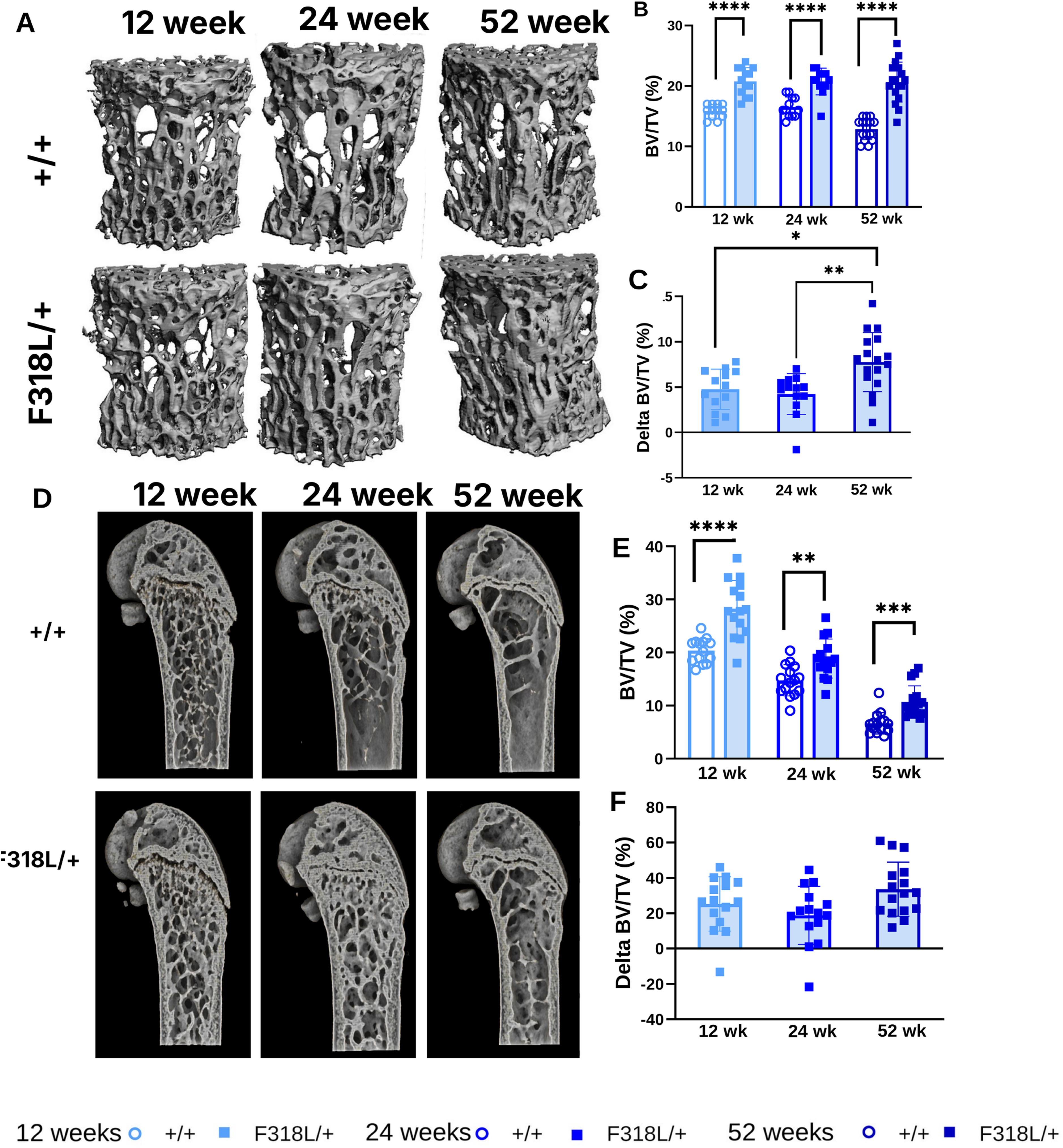
Vertebral trabecular BV/TV is increased in male F318L mouse model of ADOII. **A.** Representative 3D images of 12-, 24-, and 52-week-old male +/+ and F318L/+ vertebrae. **B**. Trabecular BV/TV from +/+ (open columns) and F318L/+ (filled columns) mice at 12-week (light blue), 24-week (dark blue), 52-week (Kelly blue) of age. **C.** Δ BV/TV at each age. **D**. Representative 3D images of 12-, 24-, and 52-week-old male +/+ and F318L/+ femurs. **E.** Trabecular BV/TV, **F**. Δ BV/TV. Data are presented as individual animals with mean ± SD. N=15/group. +/+ vs F318L/+ at each age: Mann-Whitney test; Δ across ages: Kruskal–Wallis test with Dunn’s multiple-comparisons test. *= p<0.05, **= p<0.01, ***= p<0.001, ****= p<0.0001.

### Increased femoral cortical thickness in *Clcn7* F318L/+ mice across the age span

Femurs from female F318L/+ mice demonstrated increased cortical thickness at all ages (Fig 3A, B). However, no significant impact of age was observed, as the relative increase in cortical thickness (Ct.Th) in the F318L/+ did not differ significantly across the three different age cohorts (Fig 3C). Cortical area (Ct.BA) was also increased at all ages, and cortical area/tissue area (Ct.BA/TA) at 24 and 52 weeks of age (Fig 3D, E).

**Figure 3:**
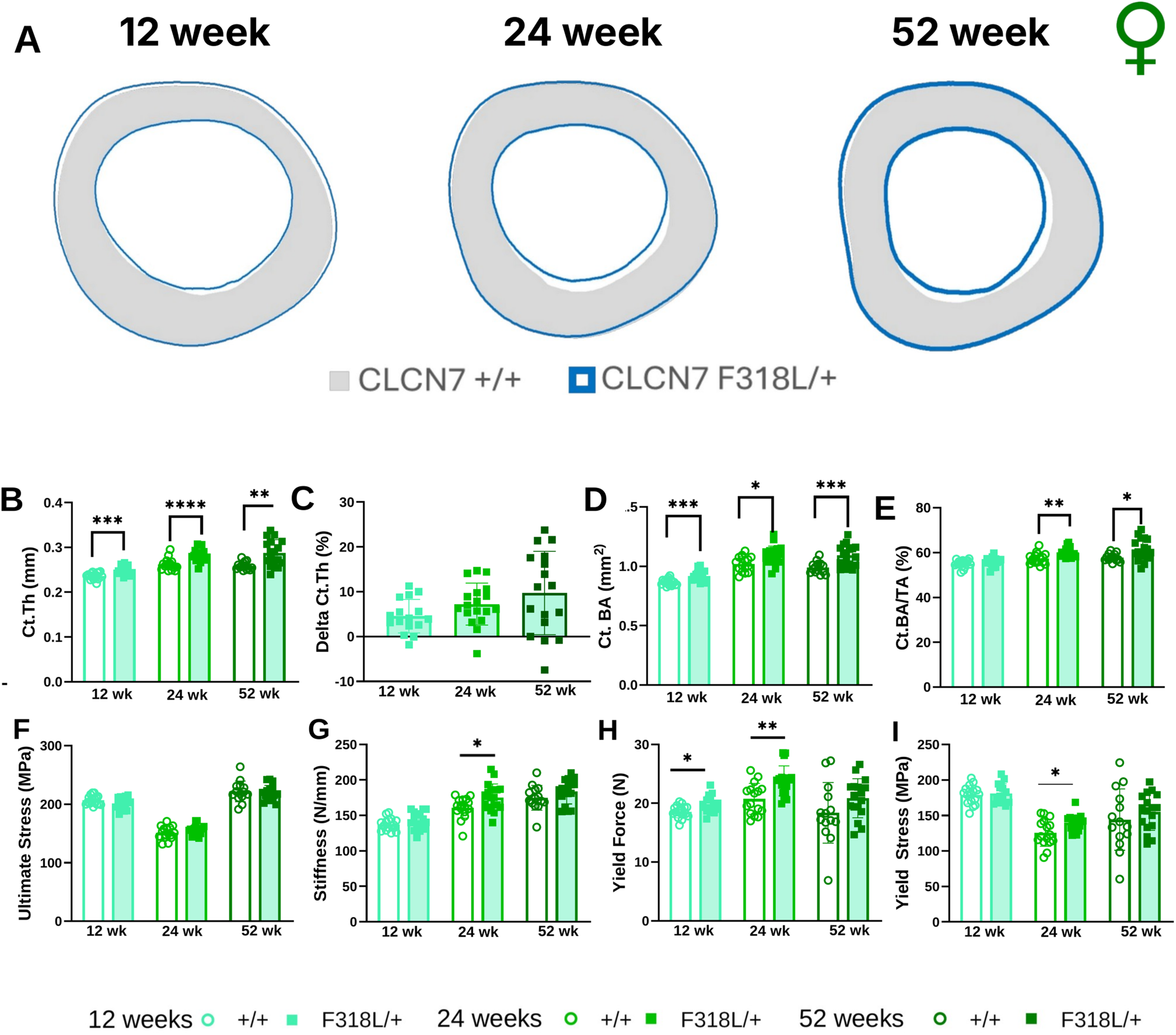
Femoral cortical bone parameters are increased in the female F318L mouse model of ADOII with no change in mechanical properties. **A.** Representative images of femoral cortical bones from 12-, 24-, and 52-week-old female +/+ and F318L/+ mice. **B. Ct.Th. C. Δ Ct.Th at each age. D–E. Cortical bone parameters, D. Ct.BA, E. Ct.BA/TA. F-I**. Femoral mechanical properties, **F**. Ultimate stress, **G**. Stiffness, **H**. Yield force, **I**. Yield stress. Data are presented as individual animals with mean ± SD. +/+ vs F318L/+ at each age: Mann-Whitney test; Δ across ages: Kruskal–Wallis test with Dunn’s multiple-comparisons test. *= p<0.05, **= p<0.01, ***= p<0.001, ****= p<0.0001.

Next, we performed 3-point bending tests to determine the femoral mechanical properties of female mice at 12, 24 and 52 weeks of age (Fig 3F–I; Supplementary Table S5). Ultimate stress did not differ between genotypes at any age (Fig 3F). Femurs from F318L/+ mice had higher yield force at 12 and 24 weeks of age (Fig 3H), and at 24 weeks of age they were also stiffer (Fig 3G) and had higher yield stress (Fig 3I) than those from +/+ littermates. At the same age, total work and post-yield displacement were lower in F318L/+ femurs (Supplementary Table S5), consistent with reduced ductility. No genotype differences were observed at 52 weeks of age.

The cortical bone phenotype in male F318L/+ mice was more modest than in females, with increased Ct.Th seen only at 12 weeks of age (Fig 4A, B), and the difference between F318L/+ and wild-type mice decreased rather than increased with age (Fig 4C). In contrast to females, 3-point bending revealed no differences in ultimate stress, stiffness, yield force or yield stress between male F318L/+ and +/+ mice at any age (Fig 4F–I), nor in any other mechanical property (Supplementary Table S5).

**Figure 4:**
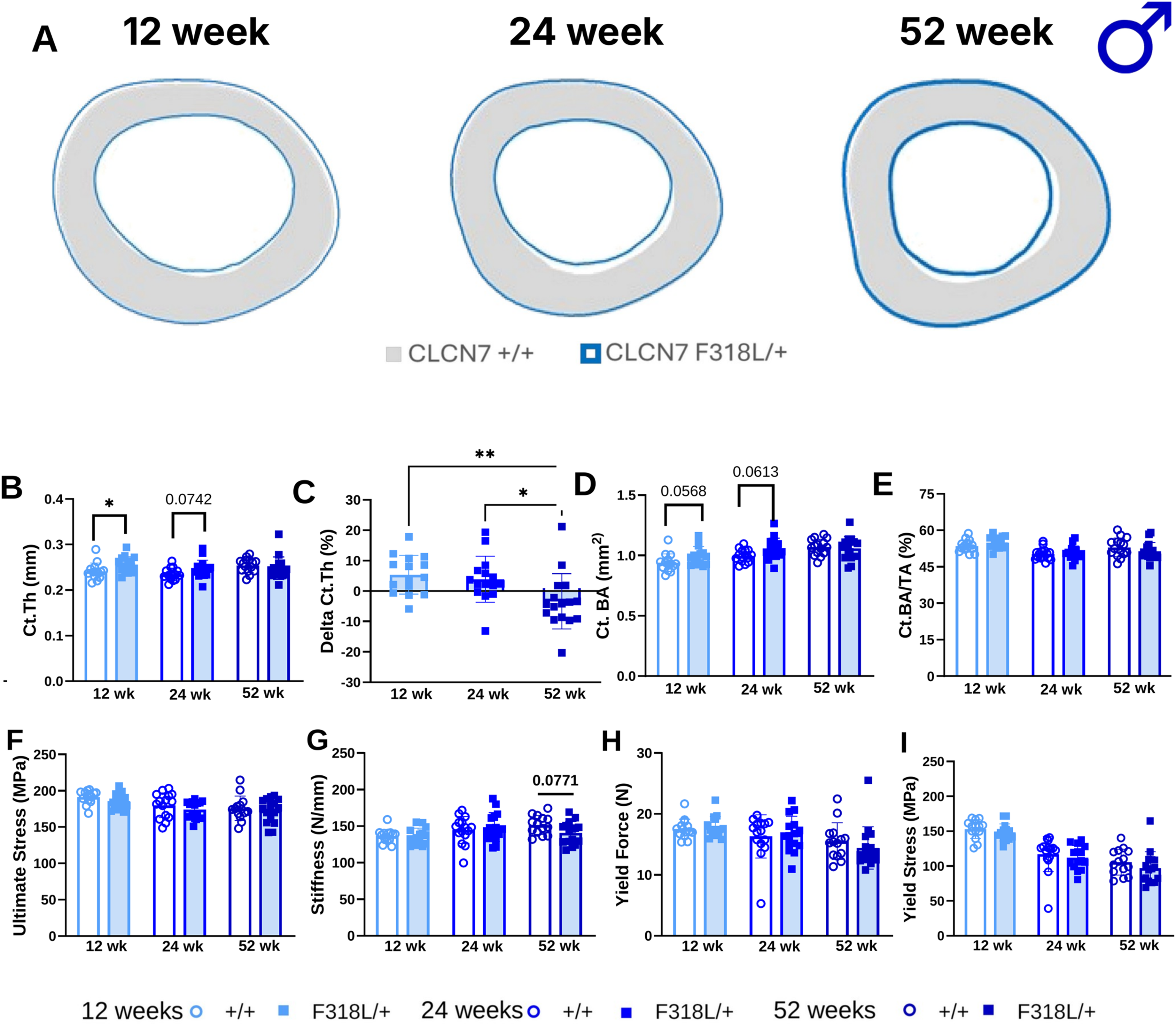
Male ADOII mice with F318L mutation have increased cortical bone with no change in mechanical properties. **A.** Representative images of femoral cortical bones from 12-, 24-, and 52-week-old male +/+ and F318L/+ mice. **B. Ct.Th. C. Δ Ct.Th at each age. D–E. Cortical bone parameters, D. Ct.BA, E. Ct.BA/TA. F-I**. Femoral mechanical properties, **F**. Ultimate stress, **G**. Stiffness, **H**. Yield force, **I**. Yield stress. Data are presented as individual animals with mean ± SD. +/+ vs F318L/+ at each age: Mann-Whitney test; Δ across ages: Kruskal–Wallis test with Dunn’s multiple-comparisons test. *= p<0.05, **= p<0.01, ***= p<0.001, ****= p<0.0001.

In summary, cortical thickness was increased at 12 weeks of age in both female and male F318L/+ mice. In females, the increase persisted at all ages and was accompanied by higher yield force at 12 and 24 weeks and by a stiffer but less ductile femur at 24 weeks of age, whereas in males the cortical phenotype waned with age and mechanical properties were unaffected. Thus, the F318L mutation affects cortical bone differently in female and male mice.

### Genetic background modifies the vertebral phenotype of ADOII mice

Given the variability in the severity of the ADOII phenotype in humans, as well as a report that mice carrying the *Clcn7* G213R mutation had more severe osteopetrosis on the 129 background than on the C57BL/6 (B6) background [37], we sought to determine whether genetic background affects the bone phenotype in ADOII mouse models. We first compared *Clcn7* F318L/+ mice on a B6 background to the previously characterized F318L/+ mice on an FVB/NJ background. At 12 weeks of age, female F318L/+ B6 mice had higher vertebral trabecular BV/TV than their +/+ littermates (20.7% vs 12.0%; Fig 5A), and the relative increase in trabecular BV/TV (ΔBV/TV) was greater on the B6 than on the FVB/NJ background (72% vs 35%; P = 0.001; Fig 5B). Male F318L/+ B6 mice likewise had higher vertebral trabecular BV/TV than their +/+ littermates (23.2% vs 16.0%; Fig 5C), but in males ΔBV/TV did not differ between the two backgrounds (45% vs 30%; P = 0.68; Fig 5D).

**Figure 5:**
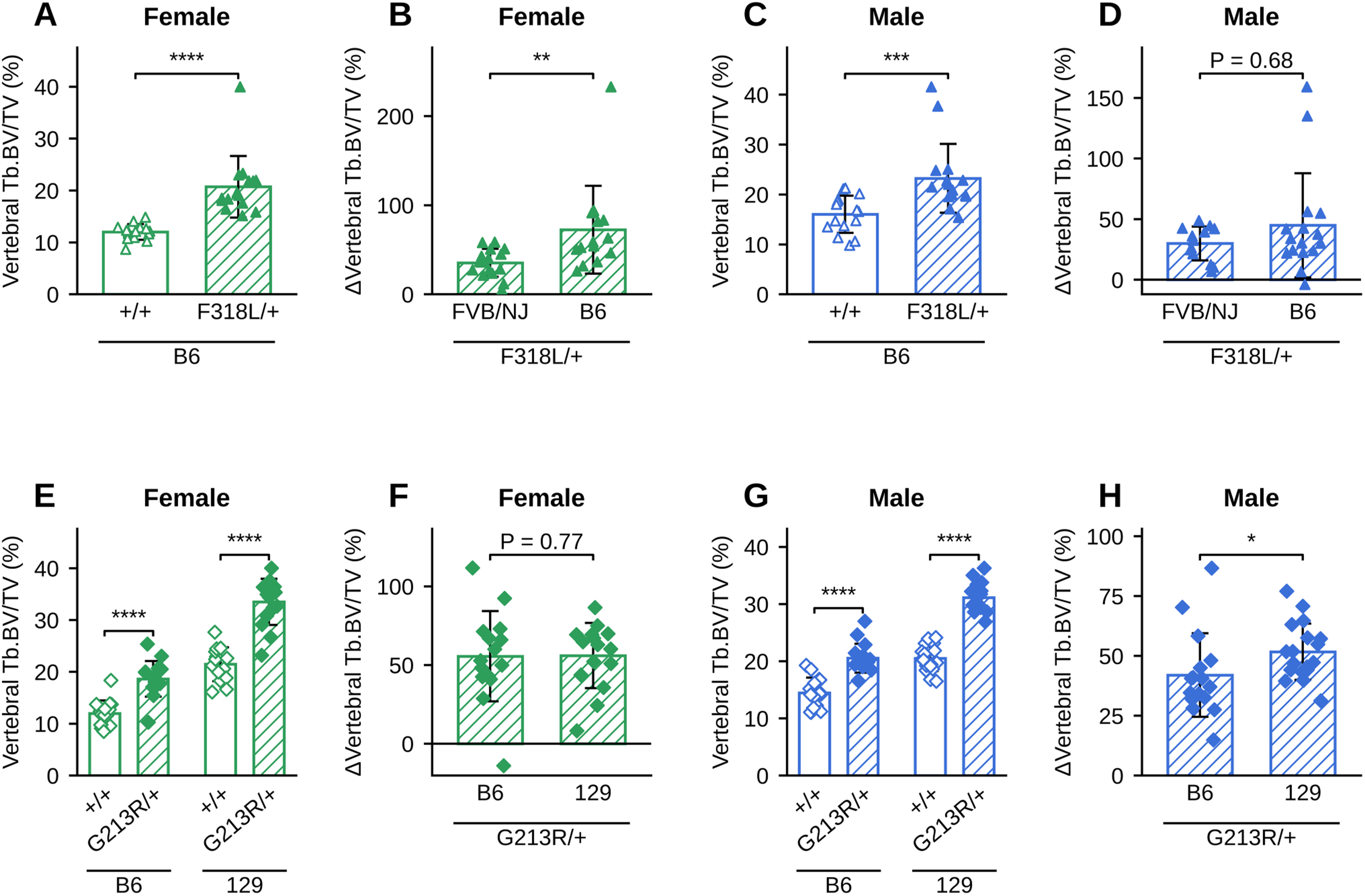
Genetic background modifies the vertebral trabecular phenotype of ADOII mice. Vertebral trabecular BV/TV of 12-week-old mice. **A–D**. F318L/+ mice. **A**. Female +/+ and F318L/+ mice on B6 background. **B**. ΔBV/TV of female F318L/+ mice on FVB/NJ and B6 backgrounds. **C, D**. As A and B, for males. **E–H**. G213R/+ mice. **E**. Female +/+ and G213R/+ mice on B6 and 129 backgrounds. **F**. ΔBV/TV of female G213R/+ mice on B6 and 129 backgrounds. **G, H**. As E and F, for males. Females are shown in green and males in blue; triangles denote F318L/+ and diamonds G213R/+ mice; open bars denote +/+ and hatched bars mutant mice. The genetic background, or the genotype shared by the compared groups, is indicated below the line under each graph. Δ (%) = (value of each mutant mouse − mean of the +/+ littermates of the same sex, line and background) / mean of those +/+ littermates × 100. Data are presented as individual animals with mean ± SD; n = 13–18/group. Mann–Whitney test. *P < 0.05, **P < 0.01, ***P < 0.001, ****P < 0.0001.

We next examined the impact of genetic background in the G213R/+ ADOII mouse model. In females, vertebral trabecular BV/TV of G213R/+ mice was increased compared to +/+ littermates on both the B6 and the 129 backgrounds (Fig 5E), and ΔBV/TV did not differ between the two backgrounds (56% vs 56%; P = 0.77; Fig 5F). Male G213R/+ mice also had higher vertebral trabecular BV/TV than +/+ littermates on both backgrounds (Fig 5G), and in males ΔBV/TV was greater on the 129 than on the B6 background (52% vs 42%; P = 0.02; Fig 5H). Because +/+ mice on the 129 background had higher baseline BV/TV (Fig 5E, G), the absolute increase in vertebral BV/TV was also larger on the 129 than on the B6 background in both females (12.0 vs 6.7 percentage points; P = 0.001) and males (10.6 vs 6.1 percentage points; P < 0.001). Thus, consistent with the previous report [37], the vertebral phenotype was more severe on the 129 background, in absolute terms in both sexes and in relative terms in males, although the difference was modest.

### Genetic background modifies the femoral trabecular and cortical phenotypes

Given the site-specific differences observed in F318L/+ mice on the FVB/NJ background (Fig 1), we next examined whether genetic background influenced the femoral phenotype. Femoral trabecular BV/TV was increased in 12-week-old female F318L/+ mice on the B6 background (15.1% vs 12.1%; Fig 6A), consistent with our findings in FVB/NJ mice (Fig 1E), but the relative increase (ΔBV/TV) was smaller on the B6 than on the FVB/NJ background (25% vs 48%; P < 0.001; Fig 6B). In males, femoral trabecular BV/TV did not differ between F318L/+ and +/+ B6 mice (Fig 6C), and ΔBV/TV was again smaller on the B6 than on the FVB/NJ background (10% vs 39%; P = 0.004; Fig 6D). Thus, the femoral trabecular phenotype of F318L/+ mice is milder on the B6 than on the FVB/NJ background. Femoral trabecular BV/TV of female G213R/+ mice was increased on both the B6 and the 129 backgrounds (Fig 6E), and ΔBV/TV did not differ between the two backgrounds (15% vs 25%; P = 0.27; Fig 6F), although the absolute increase was larger on the 129 background (4.6 vs 1.7 percentage points; P = 0.03). Likewise, male G213R/+ mice had higher femoral trabecular BV/TV than +/+ littermates on both backgrounds (Fig 6G), with similar ΔBV/TV on the two backgrounds (21% vs 27%; P = 0.20; Fig 6H).

**Figure 6:**
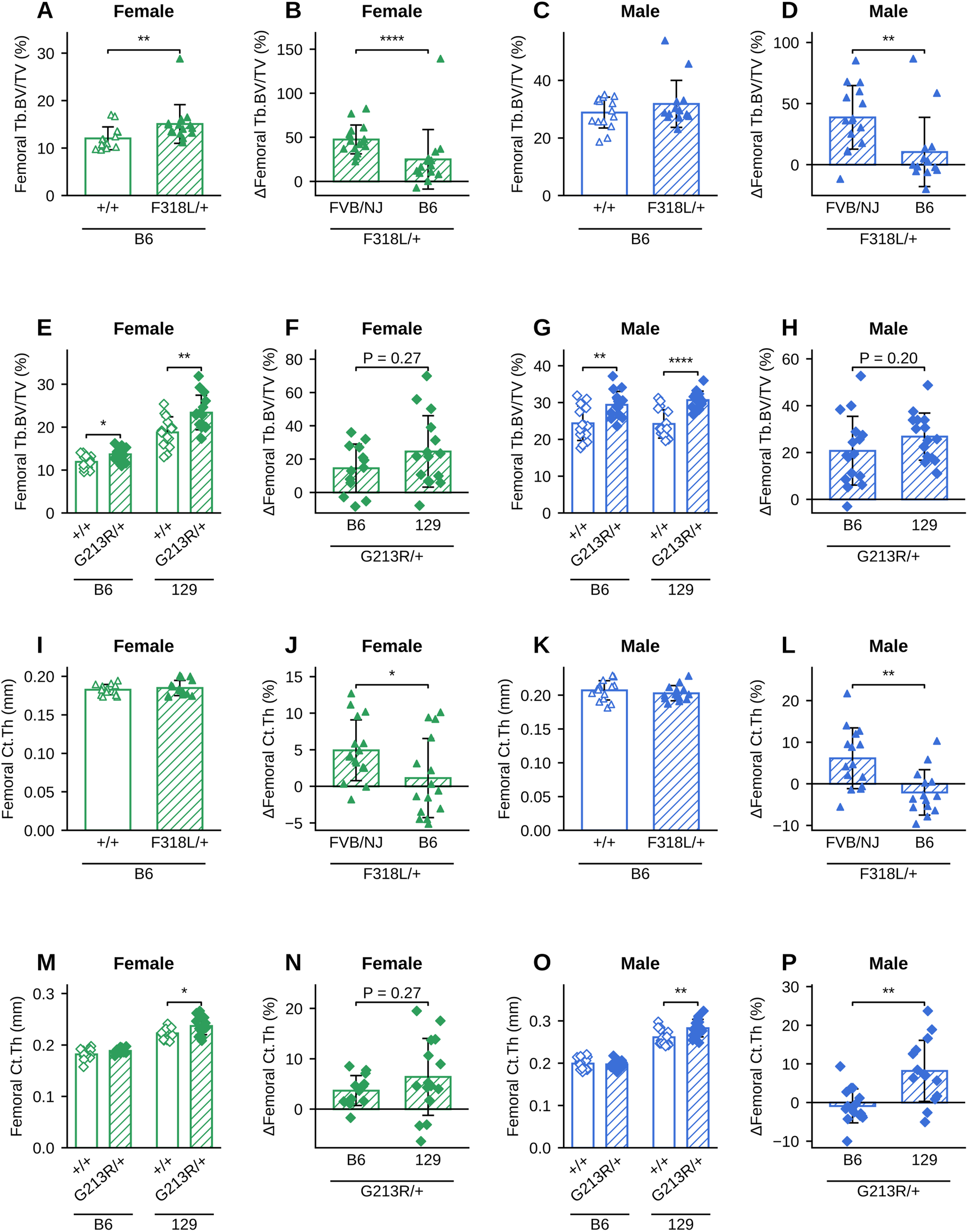
Genetic background modifies the femoral trabecular and cortical phenotypes of ADOII mice. Femoral bone of 12-week-old mice. **A–H**. Trabecular bone. **A.** Female +/+ and F318L/+ mice on B6 background. **B**. ΔBV/TV of female F318L/+ mice on FVB/NJ and B6 backgrounds. **C, D**. As A and B, for males. **E**. Female +/+ and G213R/+ mice on B6 and 129 backgrounds. **F**. ΔBV/TV of female G213R/+ mice on B6 and 129 backgrounds. **G, H**. As E and F, for males. **I–P**. Cortical thickness (Ct.Th), arranged as in A–H. **I.** Female +/+ and F318L/+ mice on B6 background. **J**. ΔCt.Th of female F318L/+ mice on FVB/NJ and B6 backgrounds. **K, L**. As I and J, for males. M. Female +/+ and G213R/+ mice on B6 and 129 backgrounds. **N**. ΔCt.Th of female G213R/+ mice on B6 and 129 backgrounds. **O, P**. As M and N, for males. Females are shown in green and males in blue; triangles denote F318L/+ and diamonds G213R/+ mice; open bars denote +/+ and hatched bars mutant mice. Data are presented as individual animals with mean ± SD; n = 13–16/group. Mann–Whitney test. *P < 0.05, **P < 0.01, ***P < 0.001, ****P < 0.0001.

We next determined whether genetic background influenced cortical bone. Cortical thickness (Ct.Th) of female F318L/+ mice on the B6 background was similar to that of +/+ littermates (Fig 6I), and the effect of the F318L mutation on Ct.Th (ΔCt.Th) was consequently smaller on the B6 than on the FVB/NJ background (1% vs 5%; P = 0.02; Fig 6J). Similarly, Ct.Th was not altered in male F318L/+ B6 mice (Fig 6K), and ΔCt.Th was smaller on the B6 than on the FVB/NJ background (−2% vs 6%; P = 0.003; Fig 6L). A similar background dependence was observed for the G213R mutation. In females, Ct.Th of G213R/+ mice was not altered on the B6 background but was increased on the 129 background (Fig 6M), although ΔCt.Th did not differ significantly between the two backgrounds (4% vs 6%; P = 0.27; Fig 6N). In males, Ct.Th of G213R/+ mice was likewise unchanged on the B6 background but increased on the 129 background (Fig 6O), and ΔCt.Th was greater on the 129 than on the B6 background (8% vs −1%; P = 0.001; Fig 6P).

### Mutation type influences the femoral trabecular phenotype only in males

Because subtle differences in disease severity between mutations are difficult to assess in human patients, we next asked whether the F318L and G213R mutations differ in their impact on the skeleton by comparing mice carrying each mutation on the same B6 background. In females, vertebral trabecular BV/TV was increased in both F318L/+ and G213R/+ mice compared to their respective +/+ littermates (Fig 7A), and ΔBV/TV did not differ between the two mutations (72% vs 56%; P = 0.49; Fig 7B). The same was true in males (Fig 7C), with similar ΔBV/TV in F318L/+ and G213R/+ mice (45% vs 42%; P = 0.36; Fig 7D). This was somewhat unexpected, given that G213R and F318L disrupt CLCN7 through distinct trafficking and functional mechanisms, respectively, but suggests that both mutations impair vertebral bone remodeling to a similar extent.

**Figure 7:**
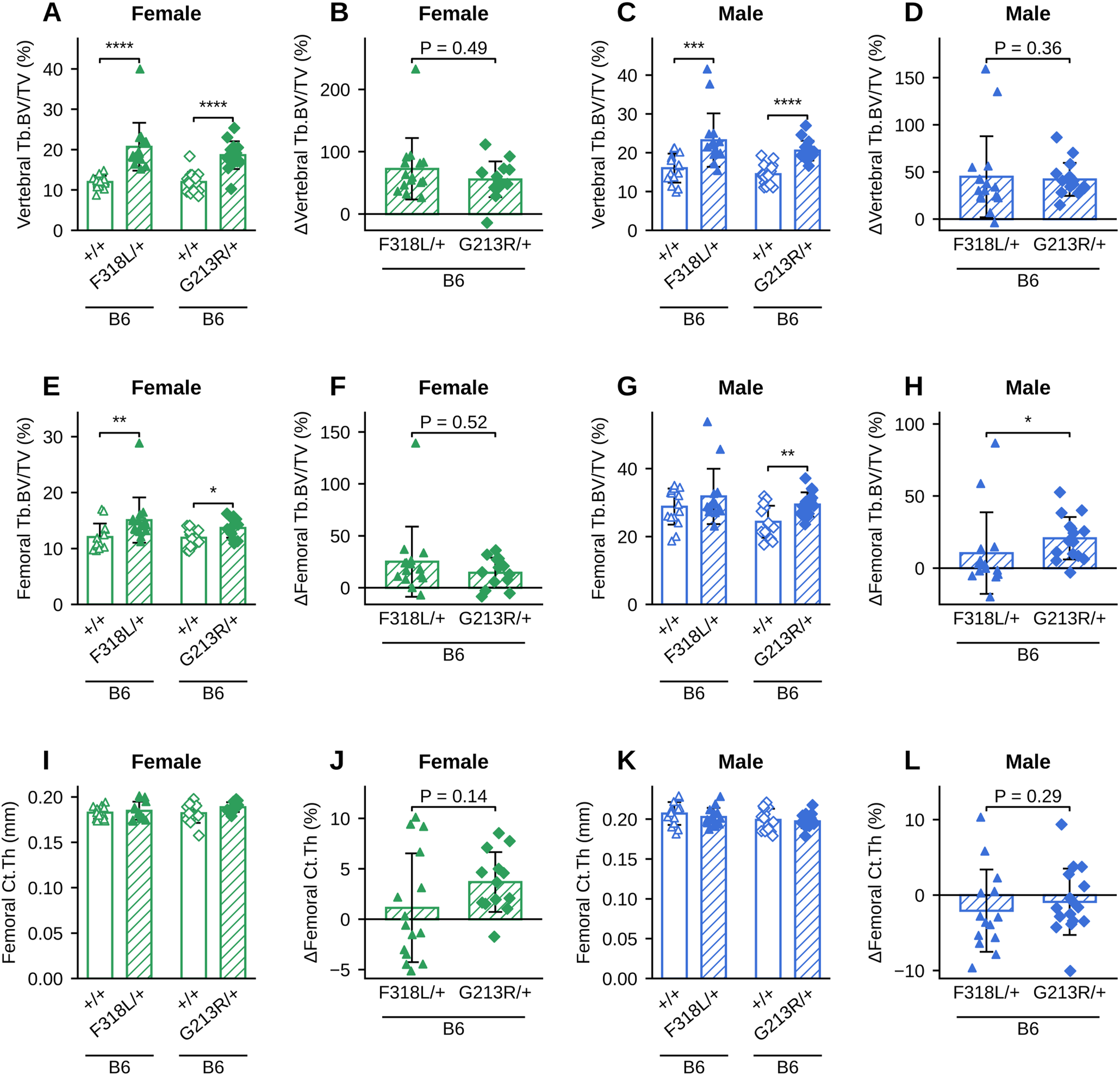
Mutation type influences the femoral trabecular phenotype in males but not the vertebral or cortical phenotypes. Comparison of 12-week-old F318L/+ and G213R/+ mice on B6 background (B6 data from Figs 5 and 6 shown again). **A–D**. Vertebral trabecular BV/TV. A. Female F318L/+ and G213R/+ mice, with corresponding +/+ littermates. B. ΔBV/TV of female F318L/+ and G213R/+ mice. **C, D**. As A and B, for males. E–H. Femoral trabecular BV/TV, arranged as in A–D. **I–L**. Femoral cortical thickness (Ct.Th), arranged as in A–D. Females are shown in green and males in blue; triangles denote F318L/+ and diamonds G213R/+ mice; open bars denote +/+ and hatched bars mutant mice. Data are presented as individual animals with mean ± SD; n = 13–16/group. Mann–Whitney test. *P < 0.05, **P < 0.01, ***P < 0.001, ****P < 0.0001.

At the femur, trabecular BV/TV was increased in female mice carrying either mutation (Fig 7E), and ΔBV/TV did not differ between the two mutations (25% vs 15%; P = 0.52; Fig 7F). In males, however, femoral trabecular BV/TV was increased in G213R/+ but not in F318L/+ mice (Fig 7G), and ΔBV/TV was greater in G213R/+ than in F318L/+ mice (21% vs 10%; P = 0.01; Fig 7H). Femoral Ct.Th was not altered by either mutation in females (Fig 7I), and ΔCt.Th did not differ between the mutations (1% vs 4%; P = 0.14; Fig 7J); likewise, in males Ct.Th was unchanged with either mutation (Fig 7K) and ΔCt.Th did not differ (−2% vs −1%; P = 0.29; Fig 7L). Thus, unlike vertebral trabecular bone, the femoral trabecular phenotype shows both background- and mutation-dependent variability, whereas cortical bone is less sensitive than trabecular bone to *Clcn7*-associated osteopetrosis and is influenced primarily by genetic background rather than mutation type.

## Discussion

In humans, clinical manifestations of autosomal dominant osteopetrosis II (ADOII) are highly variable, even when the patients share the same family tree [9]. A recent study of pediatric and adult patients with distinct *CLCN7* mutations, both with and without family history, reported greater loss of quality of life in adults compared to children, suggesting an age-dependent component of disease progression [38]. The sequelae of ADOII, especially fracture incidence, worsen with age [39]. However, whether this reflects progressive worsening of the disease or is simply due to a longer timeframe for fracture accumulation remains unclear. Furthermore, although adult human skeletons are sexually dimorphic, for example, fracture healing differs between females and males [40], limited sample sizes restrict assessment of whether there are sex differences in the ADOII phenotype.

In this study, we examined whether biological variables, including age and sex, genetic background, and specific *Clcn7* mutation, influence the osteopetrotic phenotype in murine models of ADOII. We hypothesized that these variables might modulate disease severity, possibly explaining some of the heterogeneity observed in patients. To test this, we evaluated mice across different ages, sexes, strains, and *Clcn7* mutations. Overall, trabecular bone, particularly at the vertebrae, exhibited the most robust and consistent osteopetrotic phenotype, whereas femoral cortical bone showed comparatively modest responses to *Clcn7* mutations.

To examine the effects of aging, we assessed mice carrying the F318L mutation on an FVB/NJ background at three different ages: 12 weeks old to represent the young adult group, 24 weeks old to represent the mature adult group, and 52 weeks old to represent the middle-aged to old group. At 12 weeks of age, these mice had higher trabecular bone in both vertebrae and femurs, as previously shown by the Warman group on the B6 background [20]. The relative increase in vertebral trabecular bone mass in ADOII mice compared to +/+ controls was accentuated with aging, consistent with increased clinical expression of ADOII with age.

Interestingly, the femoral trabecular phenotype was less affected by age than the vertebral phenotype, highlighting a site-specific impact of the mutation. Femoral trabecular BV/TV declined markedly between 12 and 24 weeks of age in both +/+ and F318L/+ mice (Supplementary Table S4), consistent with the early onset of age-related trabecular bone loss, which precedes cortical bone loss [26–28]. In the femur, this age-related loss may outweigh the cumulative effect of impaired osteoclast function, whereas in the vertebrae the mutant phenotype became more pronounced with age. Differences in remodeling activity between the femur and the lumbar vertebrae have been reported [41], and differences in mechanical loading between the axial and appendicular skeleton may further contribute [42, 43]. In cortical bone, the F318L mutation increased cortical thickness at all ages in females but only at 12 weeks of age in males. Despite thicker cortices, ultimate stress was unchanged. Because 3-point bending primarily probes cortical bone and the most pronounced phenotype in ADOII mice is the increase in vertebral trabecular bone, 3-point bending may not capture the most relevant differences in mechanical properties between +/+ and mutant bones; we therefore plan to test vertebral biomechanics *ex vivo* in the future.

Skeletal parameters in mice differ by sex across strains [31, 44, 45]. Osteoclasts are also sexually dimorphic [30, 46]. As such, we examined whether the ADOII phenotype varied by sex. Female and male mice exhibited comparable increases in trabecular bone mass across all ages, suggesting that sex does not substantially modulate the trabecular phenotype in ADOII. In contrast, the cortical phenotype was sexually dimorphic, with increased cortical thickness persisting with age in females, whereas in males it was limited to 12 weeks of age. Likewise, the effect of genetic background on the vertebral phenotype was observed only in females.

Baseline bone differences between strains are well-documented [33, 47]. Alam and colleagues established that genetic background influences the severity of the osteopetrotic phenotype in *Clcn7* G213R/+ mice, with phenotypic severity varying across multiple strain backgrounds. Their findings highlighted the importance of genetic modifiers in shaping disease expression [37]. We sought to determine whether a similar effect could be observed in a second ADOII model carrying the F318L mutation. Because mouse strains differ considerably in baseline trabecular bone mass, we assessed the relative effect of the mutation using ΔBV/TV compared with strain-matched wild-type controls rather than relying solely on absolute BV/TV values. We compared 12-week-old mice carrying the F318L/+ mutation in *Clcn7* on either the FVB/NJ or B6 background. The F318L/+ mutation induced markedly greater changes in femoral trabecular BV/TV and cortical thickness on the FVB/NJ than on the B6 background in both sexes, whereas its impact on vertebral BV/TV was similar in males and even greater on the B6 background in females. In contrast to the findings of Alam and colleagues [37], the relative increase in trabecular BV/TV in G213R/+ mice was only modestly greater on the 129 background, reaching significance only in the vertebrae of males, although G213R/+ increased femoral cortical thickness only on the 129 background. Site specificity of differences between strains has been reported. Prior comparisons of B6 mice to several other strains found that B6 mice had different body composition and metabolic activities resulting in lower femoral bone mass but comparable vertebral bone mass to other strains [41]. This strain-dependent effect reinforces the concept of site-specific susceptibility and may reflect differences in remodeling rates or osteoclast activity across bones.

Several factors may explain why we did not fully reproduce the genetic background effect reported by Alam and colleagues [37]. Both studies examined 12-week-old mice of both sexes and expressed the phenotype as the percent increase over strain-matched +/+ mice, but the magnitude of the phenotype differed. The direction of the background effect was preserved, but the effect size was smaller. At baseline, +/+ 129 mice had higher trabecular BV/TV than B6 mice, particularly in females (femur 19% vs 12%; vertebra 21% vs 12%), so a given absolute gain in bone resulted in a smaller relative change. The absolute increase in BV/TV was 1.7– 2.7-fold larger on the 129 than on the B6 background in the vertebrae of both sexes (P ≤ 0.001) and in the femur of females (P = 0.03), consistent with a more severe phenotype on the 129 background in absolute terms. Additionally, our mice were bred and housed at Brigham and Women’s Hospital, and genetic drift in independently maintained colonies, the precise 129 substrain, and facility-specific factors such as diet, housing and microbiota can modify skeletal phenotypes [48, 49]. Finally, differences in µCT acquisition, region-of-interest placement and segmentation thresholds influence trabecular measurements. Together, these suggest that genetic background modifies the absolute severity of the G213R/+ phenotype, though the magnitude of effect may vary.

We also examined whether different *Clcn7* mutations resulted in different phenotypes by comparing mice heterozygous for either the F318L or G213R mutation on a B6 background. As previously reported by the Econs group, who generated the model, 12-week-old G213R/+ ADOII mice had higher vertebral and femoral trabecular bone [18]. Although the G213R and F318L mutations disrupt CLCN7 function through distinct mechanisms, the relative increase in vertebral BV/TV was similar between the two models. G213R primarily causes a trafficking defect that impairs proper localization of the transporter, whereas F318L is thought to reach its normal destination but exhibits reduced Cl^-^/H^+^ exchange activity [19, 20]. Thus, we anticipated that these mechanistically distinct defects might result in different degrees of osteopetrosis. Instead, the vertebral trabecular phenotype was remarkably similar between models, suggesting that distinct mechanisms of CLCN7 dysfunction can produce comparable effects on vertebral trabecular bone. In contrast, mutation-dependent differences emerged at the femur, where G213R/+ males exhibited a greater femoral trabecular phenotype than F318L/+ males, whereas cortical thickness did not differ between the two mutations. Together, these findings indicate that the influence of mutation identity may be site-specific and that both the nature of the mutation and local skeletal factors likely contribute to phenotypic variation in ADOII.

In summary, *Clcn7* mutant mice consistently exhibited an osteopetrotic phenotype characterized by increased trabecular bone mass, and increases in cortical thickness that were more pronounced in females. The trabecular phenotype was present across age, sex, mutation, and genetic background, although age, genetic background and, in the femur, mutation type modified its magnitude in a site-specific manner. Future studies will focus on the cellular and molecular mechanisms underlying these effects and evaluate how bone mechanics and material properties contribute to the mechanical characteristics of osteopetrotic bone.

## Materials and methods

### Mice

*Clcn7* G213R/+ mice carrying a mutation analogous to CLCN7 p.G215R mutation present in the ADOII patients (referred to as *Clcn7* G213R/+) on both the C57BL/6 (B6) and 129 strain backgrounds were a generous gift of Dr. Econs [18]. B6 mice carrying the CLCN7 p.F316L mutation are analogous to the CLCN7 p.F318L mutation reported in Chinese, Italian and American ADOII patients (referred to as *Clcn7* F318L/+) were courtesy of Dr. Warman [20]. B6 Clcn7 F318L mice were backcrossed to FVB/NJ mice for 7 generations. +/+ and F318L/+ FVB/NJ littermates were co-housed and aged to 12-, 24-, and 52-week-old in our vivarium for the age comparison study. 12-week-old female and male +/+ and F318L/+ mice on the FVB/NJ and B6 backgrounds, as well as 12-week-old female and male +/+ and G213R/+ mice on the B6 and 129 backgrounds were used for strain comparison studies. 12-week-old female and male +/+, F318L/+ and G213R/+ mice on the B6 background were used for mutation comparison studies.

All mice were bred and housed at constant room temperature (22 °C ± 10%), air humidity (50% ± 20%), and a light/dark cycle of 12 h in the Brigham and Women’s Hospital/Harvard Medical School vivarium. All mice were given ad libitum access to autoclaved water and chow diet. All animal procedures were approved by the Institutional Animal Care and Use Committee, Harvard Medical School, and Brigham and Women’s Hospital.

### Micro-computed tomography (µCT)

Mice were euthanized by CO₂ inhalation, and vertebrae and femurs were harvested and stripped of soft tissue. Vertebrae were fixed in 4% PFA for 2 days before transferring to 0.4% PFA. The fourth lumbar vertebra (L4) from each mouse was scanned; when L4 was not suitable for analysis (e.g., due to vertebral fusion), L5 was analyzed instead. Vertebrae were scanned using a Scanco Medical μCT 35 machine at an isotropic voxel size of 7 μm, x-ray tube potential of 55 kVp, x-ray intensity of 0.145 mA, and an integration time of 400 ms. From the scans of the vertebrae, a region beginning 0.07 mm proximal to the growth plate and extending 1.4 mm proximally was selected for trabecular bone analysis. A semi-automated contouring approach was used to distinguish cortical and trabecular bone. The region of interest was thresholded using a global threshold that set the bone/ marrow cut off at 320 mg HA/ccm for trabecular bone. Thresholding and ROI placement were kept constant across ages, strains, and sexes; where the caudal growth plate fell within the region, the ROI was shortened to exclude it. 3-D microstructural properties of bone, including bone volume fraction (BV/TV), trabecular thickness (Tb.Th.), trabecular number (Tb.N.), and trabecular separation (Tb.Sp.) were calculated using software supplied by the manufacturer.

Femurs were collected and wrapped in PBS-soaked gauze to be kept in -20°C and were thawed for µCT analysis. µCT was performed using a Skyscan 1272 µCT system (Bruker microCT, Kontich, Belgium). Bone scans were performed using 70 kV source voltage, 142 µA current, 0.5 mm Al filter, 0.7° rotation step, 2-frame averaging, and an isotropic voxel size of 10 µm. For cortical bone analysis, a region of interest (ROI) of 0.1 mm (10 slices) centered at the femoral midshaft was used, and for trabecular bone analysis, a ROI of 1 mm in distal metaphysis extending proximally from the most proximal portion of the growth plate was used. Tissue mineral density was calculated using manufacturer supplied hydroxyapatite phantoms (0.3 and 1.25 g/cm^3^). Scans were reconstructed with NRecon (Bruker microCT) software and properly aligned using Dataviewer software (Bruker microCT). Using CT-Analyzer (CTAn; Bruker, Billerica, MA), the trabecular bone within the trabecular ROI was isolated from the cortical shell. Trabecular outcomes included bone volume fraction (BV/TV), trabecular thickness (Tb.Th), trabecular number (Tb.N), and trabecular separation (Tb.Sp). Cortical properties were calculated using a custom MATLAB (MathWorks, Natick, MA) script and included cortical thickness (Ct.Th), total cross-sectional area (TA), bone area (BA), marrow area (MA), bone area fraction (BA/TA), principal moments of inertia (Imax and Imin), and cortical tissue mineral density (TMD) [50].

### Mechanical testing on femurs

After µCT scanning, femurs were re-wrapped in PBS-soaked gauze and refrozen at −20 °C. Before mechanical testing by 3-point bending, femurs were thawed and brought to room temperature. Bones were positioned over a 7.5 mm support span, the loading point was placed at the mid-point of the bone with the anterior surface in tension, and then loaded to failure in displacement control at a rate of 0.025 mm/s. Samples were kept hydrated with PBS throughout the experiment. Load–displacement data were collected throughout testing. Geometric properties at the fracture site from µCT scans were used with standard three-point bending equations to convert force–displacement data into estimated stress–strain data. Yield was defined using the 0.2% strain offset method [50].

### Finite element analysis of vertebrae

µCT-based micro-finite element analysis (mFEA) was performed to predict the strength of the L4 vertebral body of the mice in the study using Scanco FE-software (fe_solve3, v1.16, Scanco Medical AG, Brüttisellen, Switzerland). The methods used were based on those utilized by Nyman et al. for predicting the strength of mouse vertebrae with µFEA [51]. The volume of interest for the mFEA was created by placing a 1.75 mm (250 voxels) diameter circle around the vertebral body such that it was separated from the transverse processes and posterior elements. The mFEA was performed on a 1.4 mm VOI (200 transverse slices) in the central region of the vertebral body, which did not include the vertebral body endplates. The software pre- processed the µCT images by segmenting the bone with a lower threshold of 320 mg HA/cm3, applying a Gaussian filter (sigma = 0.8, support = 1) to remove noise, removing unconnected elements with component labeling, and then converting each voxel to an 8-node brick element (7 µm × 7 µm × 7 µm) with homogeneous elastic modulus (E = 18 GPa) and Poisson’s ratio (ν = 0.3). The processed model was run to simulate high-fraction axial compression where all nodes at the caudal face of the model were constrained in all directions (x, y, and z), and the nodes at the cranial face of the model were constrained in the x- and y-directions and had a fixed negative displacement in the z-direction. The mFE model was then solved for the reaction force at the point when 2% of the model elements exceeded a von Mises equivalent strain of 7000 µε, and this force was defined as the ultimate force of the vertebral body. The apparent predicted stiffness and compression modulus of the model were also calculated.

### Statistical analysis

Data were represented in graphs as individual data points with error bars and as mean ± Standard deviation in tables. Delta values were calculated as percentage difference of the mutation groups compared to the mean of the +/+ mice. Because bone microarchitecture outcomes were measured in groups of 12–18 mice, in which normality cannot be reliably assessed, +/+ and mutant mice of the same sex, age and background, and Δ values of two groups, were compared using the two-tailed non-parametric Mann–Whitney U test, which does not assume normally distributed data or equal variances and is robust to outliers. Comparisons of three or more groups were performed with the Kruskal–Wallis test followed by Dunn’s multiple-comparisons test, its non-parametric counterpart. * p < 0.05. ** p < 0.01. *** p < 0.001, and **** p < 0.0001. To compare the magnitude of the mutant phenotype across genetic backgrounds and mutations, each parameter was expressed as the percent change of each mutant mouse relative to its wild-type littermate controls: Δ (%) = (X_mutant_ − X̄_+/+_) / X̄_+/+_ × 100, where X_mutant_ is the value of an individual mutant mouse and X̄_+/+_ is the mean value of the +/+ mice of the same sex, age, genetic background and cohort. Statistical analyses were performed using GraphPad Prism.

## Acknowledgement

Research reported in this publication was supported by NIAMS of the National Institutes of Health under grant number R01 AR078914. A total of 85% of the research was financed with federal funds. A total of 15% of the research was financed with non-governmental funds (Department of Orthopedic Surgery funds in support of JFC). We would like to acknowledge the services provided by the Center for Skeletal Research (CSR) at the Massachusetts General Hospital histology core funded by NIH P30 AR075042. The content is solely the responsibility of the authors and does not necessarily represent the official views of the National Institutes of Health.

## Data availability

The data that support the findings of this study are available from the corresponding author upon reasonable request.

---

**Supplementary Table S1.** Body weight and body length of +/+ and Clcn7-F318L/+ mice on FVB/NJ background at 12, 24, and 52 weeks of age.

| **Age (weeks)** | **Males** |  |  |  | **Females** |  |  |  |
|---|---|---|---|---|---|---|---|---|
|  | **Body weight (g)** |  | **Body length (cm)** |  | **Body weight (g)** |  | **Body length (cm)** |  |
|  | +/+ | **F318L/+** | +/+ | **F318L/+** | +/+ | **F318L/+** | +/+ | **F318L/+** |
| 12 | 28.39 ± 1.80 | 27.10 ± 1.96 | 9.70 ± 0.21 | 9.47 ± 0.28 | 21.21 ± 1.19 | 20.69 ± 1.40 | 9.10 ± 0.27 | 9.04 ± 0.25 |
| 24 | 33.70 ± 3.24 | 34.05 ± 4.56 | 9.93 ± 0.33 | 10.06 ± 0.37 | 25.69 ± 3.57 | 24.96 ± 3.22 | 9.60 ± 0.36 | 9.48 ± 0.32 |
| 52 | 36.52 ± 5.12 | 34.76 ± 4.29 | 10.22 ± 0.48 | 10.20 ± 0.37 | 27.67 ± 1.97 | 27.06 ± 3.16 | 9.70 ± 0.40 | 9.66 ± 0.27 |

**Supplementary Table S2.**
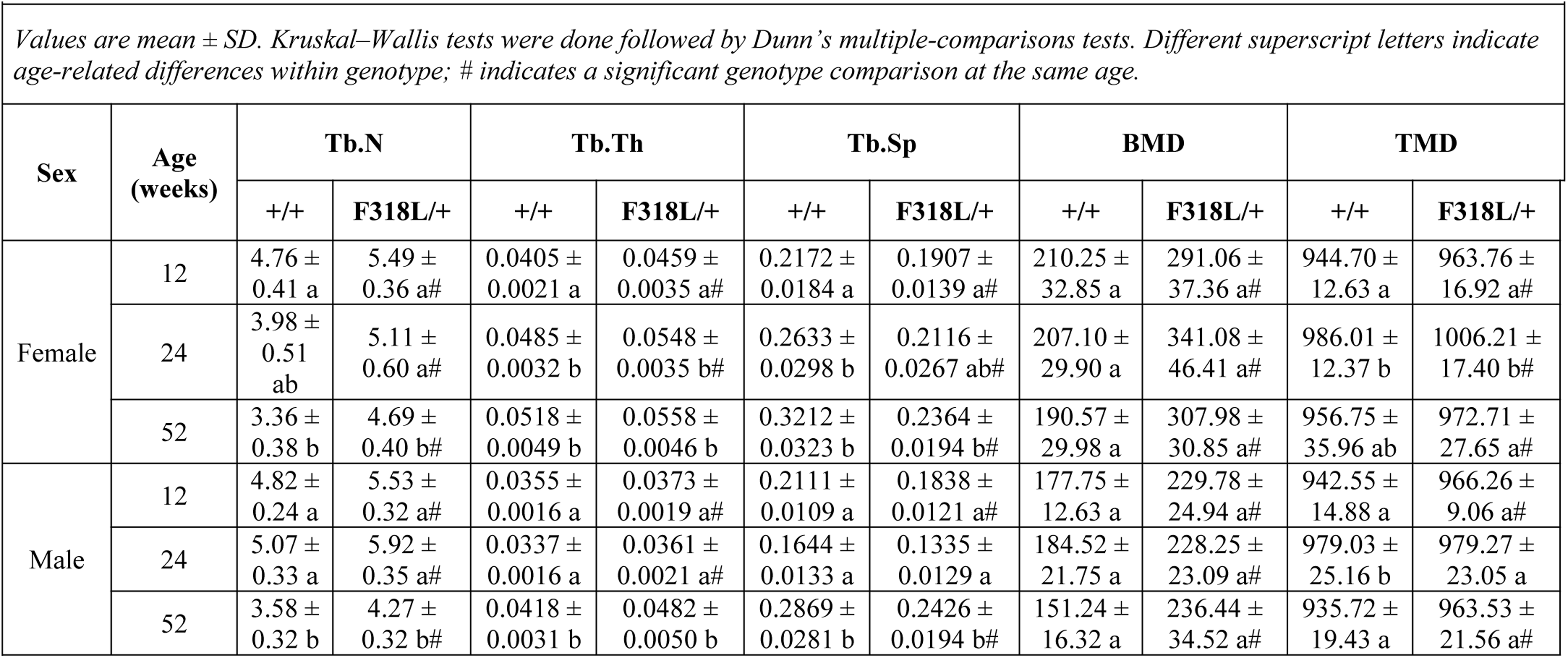
Vertebral trabecular microarchitecture in FVB/NJ +/+ and Clcn7-F318L/+ mice.

| **Sex** | **Age (weeks)** | **Tb.N** |  | **Tb.Th** |  | **Tb.Sp** |  | **BMD** |  | **TMD** |  |
|---|---|---|---|---|---|---|---|---|---|---|---|
|  |  | +/+ | **F318L/+** | +/+ | **F318L/+** | +/+ | **F318L/+** | +/+ | **F318L/+** | +/+ | **F318L/+** |
| Female | 12 | 4.76 ± 0.41 a | 5.49 ± 0.36 a# | 0.0405 ± 0.0021 a | 0.0459 ± 0.0035 a# | 0.2172 ± 0.0184 a | 0.1907 ± 0.0139 a# | 210.25 ± 32.85 a | 291.06 ± 37.36 a# | 944.70 ± 12.63 a | 963.76 ± 16.92 a# |
|  | 24 | 3.98 ± 0.51 ab | 5.11 ± 0.60 a# | 0.0485 ± 0.0032 b | 0.0548 ± 0.0035 b# | 0.2633 ± 0.0298 b | 0.2116 ± 0.0267 ab# | 207.10 ± 29.90 a | 341.08 ± 46.41 a# | 986.01 ± 12.37 b | 1006.21 ± 17.40 b# |
|  | 52 | 3.36 ± 0.38 b | 4.69 ± 0.40 b# | 0.0518 ± 0.0049 b | 0.0558 ± 0.0046 b | 0.3212 ± 0.0323 b | 0.2364 ± 0.0194 b# | 190.57 ± 29.98 a | 307.98 ± 30.85 a# | 956.75 ± 35.96 ab | 972.71 ± 27.65 a# |
| Male | 12 | 4.82 ± 0.24 a | 5.53 ± 0.32 a# | 0.0355 ± 0.0016 a | 0.0373 ± 0.0019 a# | 0.2111 ± 0.0109 a | 0.1838 ± 0.0121 a# | 177.75 ± 12.63 a | 229.78 ± 24.94 a# | 942.55 ± 14.88 a | 966.26 ± 9.06 a# |
|  | 24 | 5.07 ± 0.33 a | 5.92 ± 0.35 a# | 0.0337 ± 0.0016 a | 0.0361 ± 0.0021 a# | 0.1644 ± 0.0133 a | 0.1335 ± 0.0129 a | 184.52 ± 21.75 a | 228.25 ± 23.09 a# | 979.03 ± 25.16 b | 979.27 ± 23.05 a |
|  | 52 | 3.58 ± 0.32 b | 4.27 ± 0.32 b# | 0.0418 ± 0.0031 b | 0.0482 ± 0.0050 b | 0.2869 ± 0.0281 b | 0.2426 ± 0.0194 b# | 151.24 ± 16.32 a | 236.44 ± 34.52 a# | 935.72 ± 19.43 a | 963.53 ± 21.56 a# |

**Supplementary Table S3.** Micro-finite element (µFE)–estimated vertebral mechanics in +/+ and CLCN7 F318L/+ mice on FVB/NJ background.

| **Sex** | **Age (weeks)** | **Ultimate force (N)** |  | **Stiffness (kN/mm)** |  |
|---|---|---|---|---|---|
|  |  | **+/+** | **F318L/+** | **+/+** | **F318L/+** |
| Female | 12 | 33.56 ± 3.54 | 42.05 ± 4.13<sup>#</sup> | 4.74 ± 0.53 | 5.94 ± 0.58<sup>#</sup> |
|  | 24 | 33.61 ± 3.34 | 45.59 ± 4.44<sup>#</sup> | 4.75 ± 0.44 | 6.50 ± 0.66<sup>#</sup> |
|  | 52 | 31.61 ± 2.20 | 41.89 ± 3.06<sup>#</sup> | 4.54 ± 0.28 | 5.96 ± 0.45<sup>#</sup> |
| Male | 12 | 29.81 ± 1.35 | 34.58 ± 2.45<sup>#</sup> | 4.27 ± 0.20 | 4.91 ± 0.36<sup>#</sup> |
|  | 24 | 29.83 ± 2.25 | 34.84 ± 3.80<sup>#</sup> | 4.22 ± 0.26 | 4.96 ± 0.54<sup>#</sup> |
|  | 52 | 28.80 ± 1.77 | 36.71 ± 3.48<sup>#</sup> | 4.04 ± 0.25 | 5.19 ± 0.49<sup>#</sup> |

**Supplementary Table S4.**
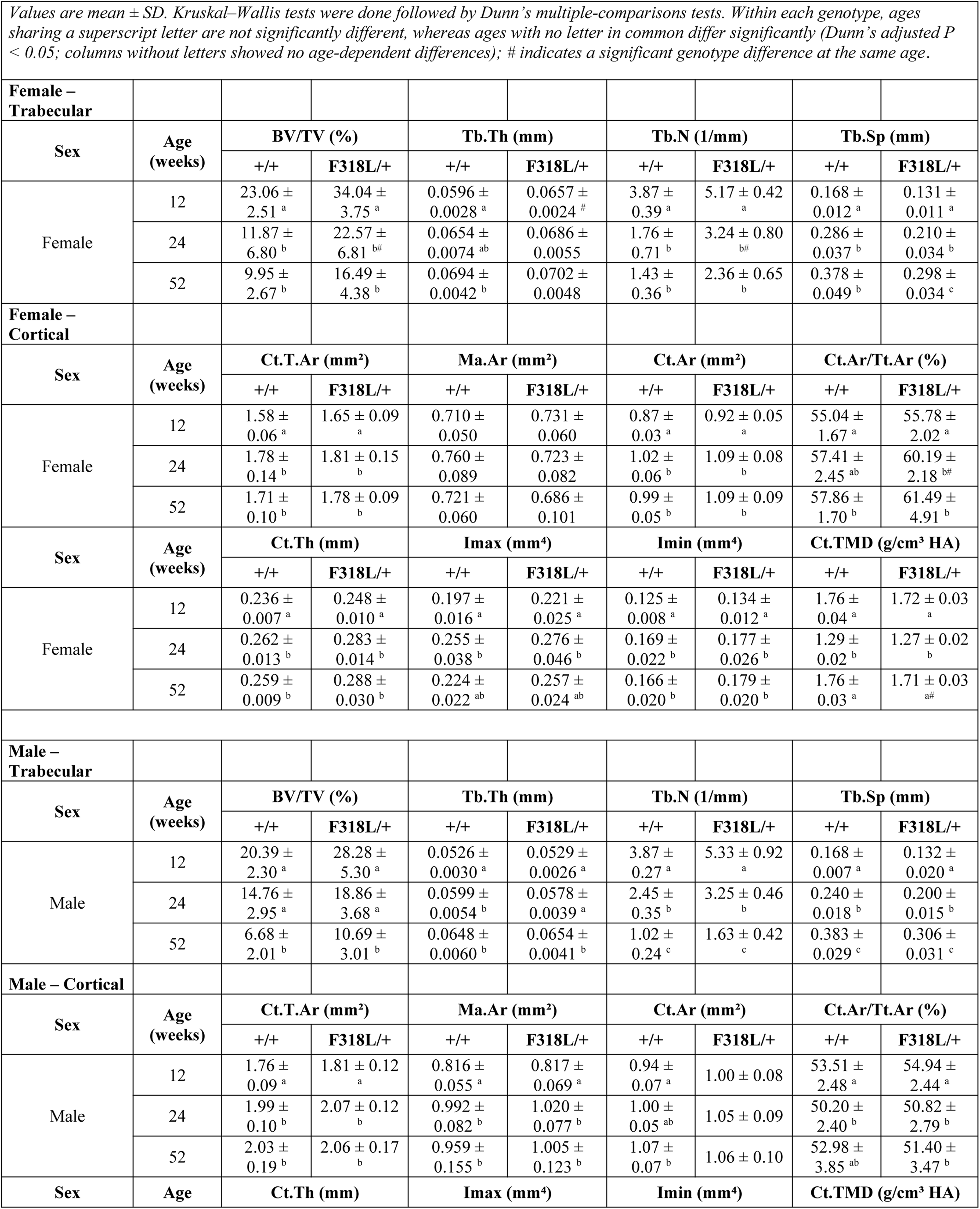
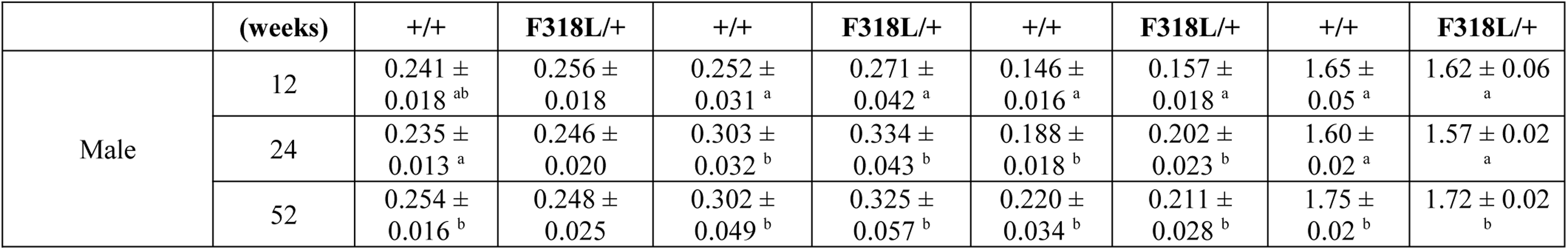
Femoral trabecular and cortical bone microarchitecture in +/+ and CLCN7 F318L/+ mice on FVB/NJ background.

| *Values are mean ± SD. Kruskal–Wallis tests were done followed by Dunn's multiple-comparisons tests. Within each genotype, ages sharing a superscript letter are not significantly different, whereas ages with no letter in common differ significantly (Dunn's adjusted P < 0.05; columns without letters showed no age-dependent differences); # indicates a significant genotype difference at the same age*. |  |  |  |  |  |  |  |  |  |
|---|---|---|---|---|---|---|---|---|---|
| **Female – Trabecular** |  |  |  |  |  |  |  |  |  |
| **Sex** | **Age (weeks)** | **BV/TV (%)** |  | **Tb.Th (mm)** |  | **Tb.N (1/mm)** |  | **Tb.Sp (mm)** |  |
|  |  | **+/+** | **F318L/+** | **+/+** | **F318L/+** | **+/+** | **F318L/+** | **+/+** | **F318L/+** |
| Female | 12 | 23.06 ± 2.51<sup>a</sup> | 34.04 ± 3.75<sup>a</sup> | 0.0596 ± 0.0028<sup>a</sup> | 0.0657 ± 0.0024<sup>#</sup> | 3.87 ± 0.39<sup>a</sup> | 5.17 ± 0.42<sup>a</sup> | 0.168 ± 0.012<sup>a</sup> | 0.131 ± 0.011<sup>a</sup> |
|  | 24 | 11.87 ± 6.80<sup>b</sup> | 22.57 ± 6.81<sup>b#</sup> | 0.0654 ± 0.0074<sup>ab</sup> | 0.0686 ± 0.0055 | 1.76 ± 0.71<sup>b</sup> | 3.24 ± 0.80<sup>b#</sup> | 0.286 ± 0.037<sup>b</sup> | 0.210 ± 0.034<sup>b</sup> |
|  | 52 | 9.95 ± 2.67<sup>b</sup> | 16.49 ± 4.38<sup>b</sup> | 0.0694 ± 0.0042<sup>b</sup> | 0.0702 ± 0.0048 | 1.43 ± 0.36<sup>b</sup> | 2.36 ± 0.65<sup>b</sup> | 0.378 ± 0.049<sup>b</sup> | 0.298 ± 0.034<sup>c</sup> |
| **Female – Cortical** |  |  |  |  |  |  |  |  |  |
| **Sex** | **Age (weeks)** | **Ct.T.Ar (mm²)** |  | **Ma.Ar (mm²)** |  | **Ct.Ar (mm²)** |  | **Ct.Ar/Tt.Ar (%)** |  |
|  |  | **+/+** | **F318L/+** | **+/+** | **F318L/+** | **+/+** | **F318L/+** | **+/+** | **F318L/+** |
| Female | 12 | 1.58 ± 0.06<sup>a</sup> | 1.65 ± 0.09<sup>a</sup> | 0.710 ± 0.050 | 0.731 ± 0.060 | 0.87 ± 0.03<sup>a</sup> | 0.92 ± 0.05<sup>a</sup> | 55.04 ± 1.67<sup>a</sup> | 55.78 ± 2.02<sup>a</sup> |
|  | 24 | 1.78 ± 0.14<sup>b</sup> | 1.81 ± 0.15<sup>b</sup> | 0.760 ± 0.089 | 0.723 ± 0.082 | 1.02 ± 0.06<sup>b</sup> | 1.09 ± 0.08<sup>b</sup> | 57.41 ± 2.45<sup>ab</sup> | 60.19 ± 2.18<sup>b#</sup> |
|  | 52 | 1.71 ± 0.10<sup>b</sup> | 1.78 ± 0.09<sup>b</sup> | 0.721 ± 0.060 | 0.686 ± 0.101 | 0.99 ± 0.05<sup>b</sup> | 1.09 ± 0.09<sup>b</sup> | 57.86 ± 1.70<sup>b</sup> | 61.49 ± 4.91<sup>b</sup> |
| **Sex** | **Age (weeks)** | **Ct.Th (mm)** |  | **Imax (mm⁴)** |  | **Imin (mm⁴)** |  | **Ct.TMD (g/cm³ HA)** |  |
|  |  | **+/+** | **F318L/+** | **+/+** | **F318L/+** | **+/+** | **F318L/+** | **+/+** | **F318L/+** |
| Female | 12 | 0.236 ± 0.007<sup>a</sup> | 0.248 ± 0.010<sup>a</sup> | 0.197 ± 0.016<sup>a</sup> | 0.221 ± 0.025<sup>a</sup> | 0.125 ± 0.008<sup>a</sup> | 0.134 ± 0.012<sup>a</sup> | 1.76 ± 0.04<sup>a</sup> | 1.72 ± 0.03<sup>a</sup> |
|  | 24 | 0.262 ± 0.013<sup>b</sup> | 0.283 ± 0.014<sup>b</sup> | 0.255 ± 0.038<sup>b</sup> | 0.276 ± 0.046<sup>b</sup> | 0.169 ± 0.022<sup>b</sup> | 0.177 ± 0.026<sup>b</sup> | 1.29 ± 0.02<sup>b</sup> | 1.27 ± 0.02<sup>b</sup> |
|  | 52 | 0.259 ± 0.009<sup>b</sup> | 0.288 ± 0.030<sup>b</sup> | 0.224 ± 0.022<sup>ab</sup> | 0.257 ± 0.024<sup>ab</sup> | 0.166 ± 0.020<sup>b</sup> | 0.179 ± 0.020<sup>b</sup> | 1.76 ± 0.03<sup>a</sup> | 1.71 ± 0.03<sup>a#</sup> |
| **Male – Trabecular** |  |  |  |  |  |  |  |  |  |
| **Sex** | **Age (weeks)** | **BV/TV (%)** |  | **Tb.Th (mm)** |  | **Tb.N (1/mm)** |  | **Tb.Sp (mm)** |  |
|  |  | **+/+** | **F318L/+** | **+/+** | **F318L/+** | **+/+** | **F318L/+** | **+/+** | **F318L/+** |
| Male | 12 | 20.39 ± 2.30<sup>a</sup> | 28.28 ± 5.30<sup>a</sup> | 0.0526 ± 0.0030<sup>a</sup> | 0.0529 ± 0.0026<sup>a</sup> | 3.87 ± 0.27<sup>a</sup> | 5.33 ± 0.92<sup>a</sup> | 0.168 ± 0.007<sup>a</sup> | 0.132 ± 0.020<sup>a</sup> |
|  | 24 | 14.76 ± 2.95<sup>a</sup> | 18.86 ± 3.68<sup>a</sup> | 0.0599 ± 0.0054<sup>b</sup> | 0.0578 ± 0.0039<sup>a</sup> | 2.45 ± 0.35<sup>b</sup> | 3.25 ± 0.46<sup>b</sup> | 0.240 ± 0.018<sup>b</sup> | 0.200 ± 0.015<sup>b</sup> |
|  | 52 | 6.68 ± 2.01<sup>b</sup> | 10.69 ± 3.01<sup>b</sup> | 0.0648 ± 0.0060<sup>b</sup> | 0.0654 ± 0.0041<sup>b</sup> | 1.02 ± 0.24<sup>c</sup> | 1.63 ± 0.42<sup>c</sup> | 0.383 ± 0.029<sup>c</sup> | 0.306 ± 0.031<sup>c</sup> |
| **Male – Cortical** |  |  |  |  |  |  |  |  |  |
| **Sex** | **Age (weeks)** | **Ct.T.Ar (mm²)** |  | **Ma.Ar (mm²)** |  | **Ct.Ar (mm²)** |  | **Ct.Ar/Tt.Ar (%)** |  |
|  |  | **+/+** | **F318L/+** | **+/+** | **F318L/+** | **+/+** | **F318L/+** | **+/+** | **F318L/+** |
| Male | 12 | 1.76 ± 0.09<sup>a</sup> | 1.81 ± 0.12<sup>a</sup> | 0.816 ± 0.055<sup>a</sup> | 0.817 ± 0.069<sup>a</sup> | 0.94 ± 0.07<sup>a</sup> | 1.00 ± 0.08 | 53.51 ± 2.48<sup>a</sup> | 54.94 ± 2.44<sup>a</sup> |
|  | 24 | 1.99 ± 0.10<sup>b</sup> | 2.07 ± 0.12<sup>b</sup> | 0.992 ± 0.082<sup>b</sup> | 1.020 ± 0.077<sup>b</sup> | 1.00 ± 0.05<sup>ab</sup> | 1.05 ± 0.09 | 50.20 ± 2.40<sup>b</sup> | 50.82 ± 2.79<sup>b</sup> |
|  | 52 | 2.03 ± 0.19<sup>b</sup> | 2.06 ± 0.17<sup>b</sup> | 0.959 ± 0.155<sup>b</sup> | 1.005 ± 0.123<sup>b</sup> | 1.07 ± 0.07<sup>b</sup> | 1.06 ± 0.10 | 52.98 ± 3.85<sup>ab</sup> | 51.40 ± 3.47<sup>b</sup> |
| **Sex** | **Age (weeks)** | **Ct.Th (mm)** |  | **Imax (mm⁴)** |  | **Imin (mm⁴)** |  | **Ct.TMD (g/cm³ HA)** |  |
|  |  | **+/+** | **F318L/+** | **+/+** | **F318L/+** | **+/+** | **F318L/+** | **+/+** | **F318L/+** |
| Male | 12 | 0.241 ± 0.018<sup>ab</sup> | 0.256 ± 0.018 | 0.252 ± 0.031<sup>a</sup> | 0.271 ± 0.042<sup>a</sup> | 0.146 ± 0.016<sup>a</sup> | 0.157 ± 0.018<sup>a</sup> | 1.65 ± 0.05<sup>a</sup> | 1.62 ± 0.06<sup>a</sup> |
|  | 24 | 0.235 ± 0.013<sup>a</sup> | 0.246 ± 0.020 | 0.303 ± 0.032<sup>b</sup> | 0.334 ± 0.043<sup>b</sup> | 0.188 ± 0.018<sup>b</sup> | 0.202 ± 0.023<sup>b</sup> | 1.60 ± 0.02<sup>a</sup> | 1.57 ± 0.02<sup>a</sup> |
|  | 52 | 0.254 ± 0.016<sup>b</sup> | 0.248 ± 0.025 | 0.302 ± 0.049<sup>b</sup> | 0.325 ± 0.057<sup>b</sup> | 0.220 ± 0.034<sup>b</sup> | 0.211 ± 0.028<sup>b</sup> | 1.75 ± 0.02<sup>b</sup> | 1.72 ± 0.02<sup>b</sup> |

**Supplementary Table S5.**
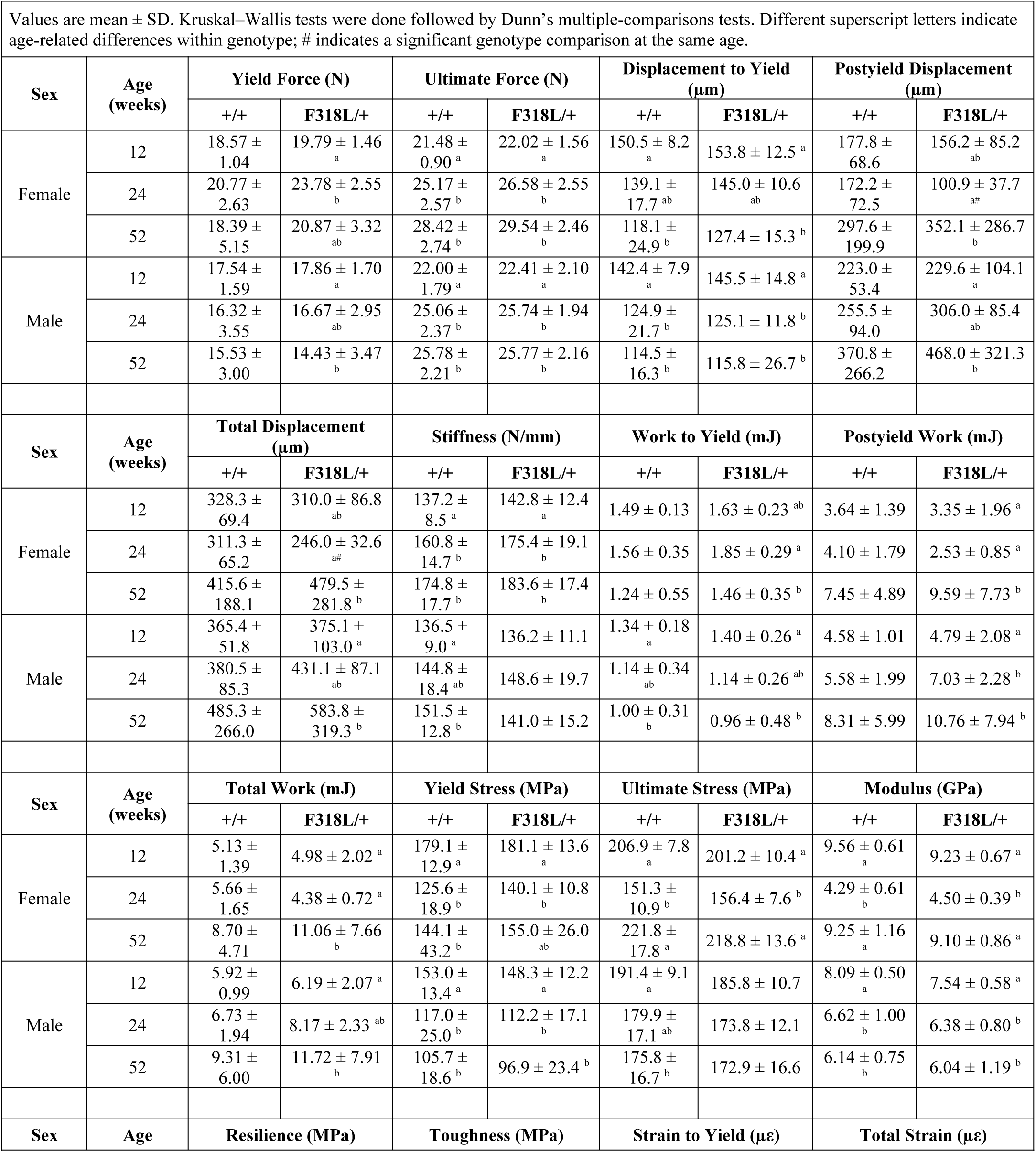
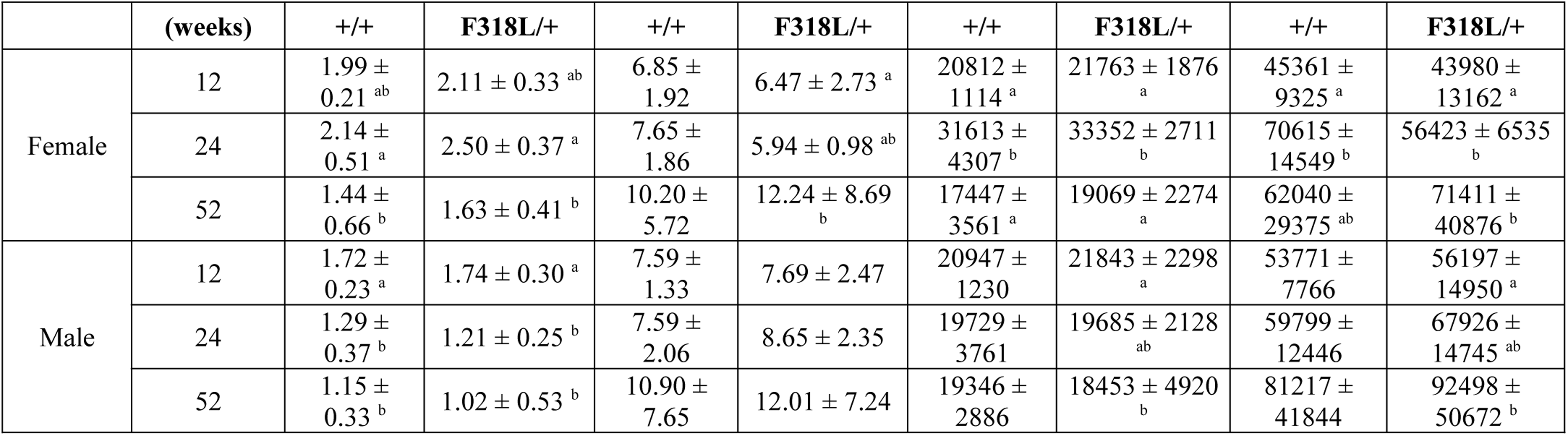
Mechanical properties of the left femur in +/+ and CLCN7 F318L/+ mice on FVB/NJ background.

| Values are mean ± SD. Kruskal–Wallis tests were done followed by Dunn’s multiple-comparisons tests. Different superscript letters indicate age-related differences within genotype; # indicates a significant genotype comparison at the same age. |  |  |  |  |  |  |  |  |  |
|---|---|---|---|---|---|---|---|---|---|
| **Sex** | **Age (weeks)** | **Yield Force (N)** |  | **Ultimate Force (N)** |  | **Displacement to Yield (µm)** |  | **Postyield Displacement (µm)** |  |
|  |  | +/+ | **F318L/+** | +/+ | **F318L/+** | +/+ | **F318L/+** | +/+ | **F318L/+** |
| Female | 12 | 18.57 ± 1.04 | 19.79 ± 1.46<sup>a</sup> | 21.48 ± 0.90<sup>a</sup> | 22.02 ± 1.56<sup>a</sup> | 150.5 ± 8.2<sup>a</sup> | 153.8 ± 12.5<sup>a</sup> | 177.8 ± 68.6 | 156.2 ± 85.2<sup>ab</sup> |
|  | 24 | 20.77 ± 2.63 | 23.78 ± 2.55<sup>b</sup> | 25.17 ± 2.57<sup>b</sup> | 26.58 ± 2.55<sup>b</sup> | 139.1 ± 17.7<sup>ab</sup> | 145.0 ± 10.6<sup>ab</sup> | 172.2 ± 72.5 | 100.9 ± 37.7<sup>a#</sup> |
|  | 52 | 18.39 ± 5.15 | 20.87 ± 3.32<sup>ab</sup> | 28.42 ± 2.74<sup>b</sup> | 29.54 ± 2.46<sup>b</sup> | 118.1 ± 24.9<sup>b</sup> | 127.4 ± 15.3<sup>b</sup> | 297.6 ± 199.9 | 352.1 ± 286.7<sup>b</sup> |
| Male | 12 | 17.54 ± 1.59 | 17.86 ± 1.70<sup>a</sup> | 22.00 ± 1.79<sup>a</sup> | 22.41 ± 2.10<sup>a</sup> | 142.4 ± 7.9<sup>a</sup> | 145.5 ± 14.8<sup>a</sup> | 223.0 ± 53.4 | 229.6 ± 104.1<sup>a</sup> |
|  | 24 | 16.32 ± 3.55 | 16.67 ± 2.95<sup>ab</sup> | 25.06 ± 2.37<sup>b</sup> | 25.74 ± 1.94<sup>b</sup> | 124.9 ± 21.7<sup>b</sup> | 125.1 ± 11.8<sup>b</sup> | 255.5 ± 94.0 | 306.0 ± 85.4<sup>ab</sup> |
|  | 52 | 15.53 ± 3.00 | 14.43 ± 3.47<sup>b</sup> | 25.78 ± 2.21<sup>b</sup> | 25.77 ± 2.16<sup>b</sup> | 114.5 ± 16.3<sup>b</sup> | 115.8 ± 26.7<sup>b</sup> | 370.8 ± 266.2 | 468.0 ± 321.3<sup>b</sup> |
| **Sex** | **Age (weeks)** | **Total Displacement (µm)** |  | **Stiffness (N/mm)** |  | **Work to Yield (mJ)** |  | **Postyield Work (mJ)** |  |
|  |  | +/+ | **F318L/+** | +/+ | **F318L/+** | +/+ | **F318L/+** | +/+ | **F318L/+** |
| Female | 12 | 328.3 ± 69.4 | 310.0 ± 86.8<sup>ab</sup> | 137.2 ± 8.5<sup>a</sup> | 142.8 ± 12.4<sup>a</sup> | 1.49 ± 0.13 | 1.63 ± 0.23<sup>ab</sup> | 3.64 ± 1.39 | 3.35 ± 1.96<sup>a</sup> |
|  | 24 | 311.3 ± 65.2 | 246.0 ± 32.6<sup>a#</sup> | 160.8 ± 14.7<sup>b</sup> | 175.4 ± 19.1<sup>b</sup> | 1.56 ± 0.35 | 1.85 ± 0.29<sup>a</sup> | 4.10 ± 1.79 | 2.53 ± 0.85<sup>a</sup> |
|  | 52 | 415.6 ± 188.1 | 479.5 ± 281.8<sup>b</sup> | 174.8 ± 17.7<sup>b</sup> | 183.6 ± 17.4<sup>b</sup> | 1.24 ± 0.55 | 1.46 ± 0.35<sup>b</sup> | 7.45 ± 4.89 | 9.59 ± 7.73<sup>b</sup> |
| Male | 12 | 365.4 ± 51.8 | 375.1 ± 103.0<sup>a</sup> | 136.5 ± 9.0<sup>a</sup> | 136.2 ± 11.1 | 1.34 ± 0.18<sup>a</sup> | 1.40 ± 0.26<sup>a</sup> | 4.58 ± 1.01 | 4.79 ± 2.08<sup>a</sup> |
|  | 24 | 380.5 ± 85.3 | 431.1 ± 87.1<sup>ab</sup> | 144.8 ± 18.4<sup>ab</sup> | 148.6 ± 19.7 | 1.14 ± 0.34<sup>ab</sup> | 1.14 ± 0.26<sup>ab</sup> | 5.58 ± 1.99 | 7.03 ± 2.28<sup>b</sup> |
|  | 52 | 485.3 ± 266.0 | 583.8 ± 319.3<sup>b</sup> | 151.5 ± 12.8<sup>b</sup> | 141.0 ± 15.2 | 1.00 ± 0.31<sup>b</sup> | 0.96 ± 0.48<sup>b</sup> | 8.31 ± 5.99 | 10.76 ± 7.94<sup>b</sup> |
| **Sex** | **Age (weeks)** | **Total Work (mJ)** |  | **Yield Stress (MPa)** |  | **Ultimate Stress (MPa)** |  | **Modulus (GPa)** |  |
|  |  | +/+ | **F318L/+** | +/+ | **F318L/+** | +/+ | **F318L/+** | +/+ | **F318L/+** |
| Female | 12 | 5.13 ± 1.39 | 4.98 ± 2.02<sup>a</sup> | 179.1 ± 12.9<sup>a</sup> | 181.1 ± 13.6<sup>a</sup> | 206.9 ± 7.8<sup>a</sup> | 201.2 ± 10.4<sup>a</sup> | 9.56 ± 0.61<sup>a</sup> | 9.23 ± 0.67<sup>a</sup> |
|  | 24 | 5.66 ± 1.65 | 4.38 ± 0.72<sup>a</sup> | 125.6 ± 18.9<sup>b</sup> | 140.1 ± 10.8<sup>b</sup> | 151.3 ± 10.9<sup>b</sup> | 156.4 ± 7.6<sup>b</sup> | 4.29 ± 0.61<sup>b</sup> | 4.50 ± 0.39<sup>b</sup> |
|  | 52 | 8.70 ± 4.71 | 11.06 ± 7.66<sup>b</sup> | 144.1 ± 43.2<sup>b</sup> | 155.0 ± 26.0<sup>ab</sup> | 221.8 ± 17.8<sup>a</sup> | 218.8 ± 13.6<sup>a</sup> | 9.25 ± 1.16<sup>a</sup> | 9.10 ± 0.86<sup>a</sup> |
| Male | 12 | 5.92 ± 0.99 | 6.19 ± 2.07<sup>a</sup> | 153.0 ± 13.4<sup>a</sup> | 148.3 ± 12.2<sup>a</sup> | 191.4 ± 9.1<sup>a</sup> | 185.8 ± 10.7 | 8.09 ± 0.50<sup>a</sup> | 7.54 ± 0.58<sup>a</sup> |
|  | 24 | 6.73 ± 1.94 | 8.17 ± 2.33<sup>ab</sup> | 117.0 ± 25.0<sup>b</sup> | 112.2 ± 17.1<sup>b</sup> | 179.9 ± 17.1<sup>ab</sup> | 173.8 ± 12.1 | 6.62 ± 1.00<sup>b</sup> | 6.38 ± 0.80<sup>b</sup> |
|  | 52 | 9.31 ± 6.00 | 11.72 ± 7.91<sup>b</sup> | 105.7 ± 18.6<sup>b</sup> | 96.9 ± 23.4<sup>b</sup> | 175.8 ± 16.7<sup>b</sup> | 172.9 ± 16.6 | 6.14 ± 0.75<sup>b</sup> | 6.04 ± 1.19<sup>b</sup> |
| **Sex** | **Age** | **Resilience (MPa)** |  | **Toughness (MPa)** |  | **Strain to Yield (µε)** |  | **Total Strain (µε)** |  |
|  | **(weeks)** | +/+ | **F318L/+** | +/+ | **F318L/+** | +/+ | **F318L/+** | +/+ | **F318L/+** |
| Female | 12 | 1.99 ± 0.21<sup>ab</sup> | 2.11 ± 0.33<sup>ab</sup> | 6.85 ± 1.92 | 6.47 ± 2.73<sup>a</sup> | 20812 ± 1114<sup>a</sup> | 21763 ± 1876<sup>a</sup> | 45361 ± 9325<sup>a</sup> | 43980 ± 13162<sup>a</sup> |
|  | 24 | 2.14 ± 0.51<sup>a</sup> | 2.50 ± 0.37<sup>a</sup> | 7.65 ± 1.86 | 5.94 ± 0.98<sup>ab</sup> | 31613 ± 4307<sup>b</sup> | 33352 ± 2711<sup>b</sup> | 70615 ± 14549<sup>b</sup> | 56423 ± 6535<sup>b</sup> |
|  | 52 | 1.44 ± 0.66<sup>b</sup> | 1.63 ± 0.41<sup>b</sup> | 10.20 ± 5.72 | 12.24 ± 8.69<sup>b</sup> | 17447 ± 3561<sup>a</sup> | 19069 ± 2274<sup>a</sup> | 62040 ± 29375<sup>ab</sup> | 71411 ± 40876<sup>b</sup> |
| Male | 12 | 1.72 ± 0.23<sup>a</sup> | 1.74 ± 0.30<sup>a</sup> | 7.59 ± 1.33 | 7.69 ± 2.47 | 20947 ± 1230 | 21843 ± 2298<sup>a</sup> | 53771 ± 7766 | 56197 ± 14950<sup>a</sup> |
|  | 24 | 1.29 ± 0.37<sup>b</sup> | 1.21 ± 0.25<sup>b</sup> | 7.59 ± 2.06 | 8.65 ± 2.35 | 19729 ± 3761 | 19685 ± 2128<sup>ab</sup> | 59799 ± 12446 | 67926 ± 14745<sup>ab</sup> |
|  | 52 | 1.15 ± 0.33<sup>b</sup> | 1.02 ± 0.53<sup>b</sup> | 10.90 ± 7.65 | 12.01 ± 7.24 | 19346 ± 2886 | 18453 ± 4920<sup>b</sup> | 81217 ± 41844 | 92498 ± 50672<sup>b</sup> |

## References

1. Wu, C.C., et al., Diagnosis and Management of Osteopetrosis: Consensus Guidelines From the Osteopetrosis Working Group. J Clin Endocrinol Metab, 2017. 102(9): p. 3111–3123.

2. Penna, S., et al., One Disease, Many Genes: Implications for the Treatment of Osteopetroses. Front Endocrinol (Lausanne), 2019. 10: p. 85.

3. Palagano, E., et al., Genetics of Osteopetrosis. Curr Osteoporos Rep, 2018. 16(1): p. 13–25.

4. Waguespack, S.G., et al., Chloride channel 7 (ClCN7) gene mutations and autosomal dominant osteopetrosis, type II. J Bone Miner Res, 2003. 18(8): p. 1513–8.

5. Cleiren, E., et al., Albers-Schönberg disease (autosomal dominant osteopetrosis, type II) results from mutations in the ClCN7 chloride channel gene. Hum Mol Genet, 2001. 10(25): p. 2861–7.

6. Bollerslev, J., et al., Autosomal dominant osteopetrosis revisited: lessons from recent studies. Eur J Endocrinol, 2013. 169(2): p. R39–57.

7. Wada, K., et al., A case of autosomal dominant osteopetrosis type II with a novel TCIRG1 gene mutation. J Pediatr Endocrinol Metab, 2013. 26(5-6): p. 575–7.

8. Jodeh, W., et al., Autosomal Dominant Osteopetrosis (ADO) Caused by a Missense Variant in the TCIRG1 Gene. J Clin Endocrinol Metab, 2024. 109(7): p. 1726–1732.

9. Polgreen, L.E., E.A. Imel, and M.J. Econs, Autosomal dominant osteopetrosis. Bone, 2023. 170: p. 116723.

10. Del Fattore, A., et al., Clinical, genetic, and cellular analysis of 49 osteopetrotic patients: implications for diagnosis and treatment. J Med Genet, 2006. 43(4): p. 315–25.

11. Bénichou, O.D., J.D. Laredo, and M.C. de Vernejoul, Type II autosomal dominant osteopetrosis (Albers-Schönberg disease): clinical and radiological manifestations in 42 patients. Bone, 2000. 26(1): p. 87–93.

12. Bollerslev, J., et al., Structural and histomorphometric studies of iliac crest trabecular and cortical bone in autosomal dominant osteopetrosis: a study of two radiological types. Bone, 1989. 10(1): p. 19–24.

13. Ludwig, C.F., et al., Common gating of both CLC transporter subunits underlies voltage-dependent activation of the 2Cl-/1H+ exchanger ClC-7/Ostm1. J Biol Chem, 2013. 288(40): p. 28611–9.

14. Brandt, S. and T.J. Jentsch, ClC-6 and ClC-7 are two novel broadly expressed members of the CLC chloride channel family. FEBS Lett, 1995. 377(1): p. 15–20.

15. Zifarelli, G., The Role of the Lysosomal Cl(-)/H(+) Antiporter ClC-7 in Osteopetrosis and Neurodegeneration. Cells, 2022. 11(3).

16. Kornak, U., et al., Loss of the ClC-7 chloride channel leads to osteopetrosis in mice and man. Cell, 2001. 104(2): p. 205–15.

17. Funck-Brentano, T., et al., Osteopetrosis and related osteoclast disorders in adults: A review and knowledge gaps On behalf of the European calcified tissue society and ERN BOND. Eur J Med Genet, 2024. 69: p. 104936.

18. Alam, I., et al., Generation of the first autosomal dominant osteopetrosis type II (ADO2) disease models. Bone, 2014. 59: p. 66–75.

19. Schulz, P., et al., The G215R mutation in the Cl-/H+-antiporter ClC-7 found in ADO II osteopetrosis does not abolish function but causes a severe trafficking defect. PLoS One, 2010. 5(9): p. e12585.

20. Caetano-Lopes, J., et al., Clcn7(F318L/+) as a new mouse model of Albers-Schönberg disease. Bone, 2017. 105: p. 253–261.

21. Zaidi, M., Skeletal remodeling in health and disease. Nat Med, 2007. 13(7): p. 791–801.

22. Havill, L.M., et al., Effects of genes, sex, age, and activity on BMC, bone size, and areal and volumetric BMD. J Bone Miner Res, 2007. 22(5): p. 737–46.

23. Plotkin, L.I., A. Bruzzaniti, and R. Pianeta, Sexual Dimorphism in the Musculoskeletal System: Sex Hormones and Beyond. J Endocr Soc, 2024. 8(10): p. bvae153.

24. Welc, S.S., et al., Aging: A struggle for beneficial to overcome negative factors made by muscle and bone. Mech Ageing Dev, 2025. 224: p. 112039.

25. da Costa, J.C., et al., Physical Activity and Sedentary Behavior on Bone Health in Adults: A Systematic Review. Int J Sports Med, 2025. 46(3): p. 164–171.

26. Bonewald, L., Use it or lose it to age: A review of bone and muscle communication. Bone, 2019. 120: p. 212–218.

27. Khosla, S., Pathogenesis of age-related bone loss in humans. J Gerontol A Biol Sci Med Sci, 2013. 68(10): p. 1226–35.

28. Riggs, B.L., et al., Population-based study of age and sex differences in bone volumetric density, size, geometry, and structure at different skeletal sites. J Bone Miner Res, 2004. 19(12): p. 1945–54.

29. Alswat, K.A., Gender Disparities in Osteoporosis. J Clin Med Res, 2017. 9(5): p. 382–387.

30. Lorenzo, J., Sexual Dimorphism in Osteoclasts. Cells, 2020. 9(9).

31. Lu, D., et al., Temporal and Quantitative Transcriptomic Differences Define Sexual Dimorphism in Murine Postnatal Bone Aging. JBMR Plus, 2022. 6(2): p. e10579.

32. Orwoll, E.S., J.K. Belknap, and R.F. Klein, Gender specificity in the genetic determinants of peak bone mass. J Bone Miner Res, 2001. 16(11): p. 1962–71.

33. Papageorgiou, M., et al., Age- and Strain-Related Differences in Bone Microstructure and Body Composition During Development in Inbred Male Mouse Strains. Calcif Tissue Int, 2020. 106(4): p. 431–443.

34. Beamer, W.G., et al., Genetic variability in adult bone density among inbred strains of mice. Bone, 1996. 18(5): p. 397–403.

35. Sabsovich, I., et al., Bone microstructure and its associated genetic variability in 12 inbred mouse strains: microCT study and in silico genome scan. Bone, 2008. 42(2): p. 439–51.

36. Voide, R., G.H. van Lenthe, and R. Müller, Bone morphometry strongly predicts cortical bone stiffness and strength, but not toughness, in inbred mouse models of high and low bone mass. J Bone Miner Res, 2008. 23(8): p. 1194–203.

37. Alam, I., et al., Phenotypic severity of autosomal dominant osteopetrosis type II (ADO2) mice on different genetic backgrounds recapitulates the features of human disease. Bone, 2017. 94: p. 34–41.

38. Polgreen, L.E., et al., Patient-Reported Outcomes in Autosomal Dominant Osteopetrosis: Findings From the Osteopetrosis Registry Study. J Clin Endocrinol Metab, 2025. 110(3): p. e607–e614.

39. Waguespack, S.G., et al., Autosomal dominant osteopetrosis: clinical severity and natural history of 94 subjects with a chloride channel 7 gene mutation. J Clin Endocrinol Metab, 2007. 92(3): p. 771–8.

40. Kurapaty, S.S. and W.K. Hsu, Sex-Based Difference in Bone Healing: A Review of Recent Pre- clinical Literature. Curr Rev Musculoskelet Med, 2022. 15(6): p. 651–658.

41. Kim, M.Y., et al., Metabolic activities affect femur and lumbar vertebrae remodeling, and anti-resorptive risedronate disturbs femoral cortical bone remodeling. Exp Mol Med, 2021. 53(1): p. 103–114.

42. Skerry, T.M., The response of bone to mechanical loading and disuse: fundamental principles and influences on osteoblast/osteocyte homeostasis. Arch Biochem Biophys, 2008. 473(2): p. 117–23.

43. Bikle, D.D. and B.P. Halloran, The response of bone to unloading. J Bone Miner Metab, 1999. 17(4): p. 233–44.

44. Glatt, V., et al., Age-related changes in trabecular architecture differ in female and male C57BL/6J mice. J Bone Miner Res, 2007. 22(8): p. 1197–207.

45. Kerschan-Schindl, K., et al., Assessment of Bone Microstructure by Micro CT in C57BL/6J Mice for Sex-Specific Differentiation. Int J Mol Sci, 2022. 23(23).

46. Mun, S.H., et al., Sexual Dimorphism in Differentiating Osteoclast Precursors Demonstrates Enhanced Inflammatory Pathway Activation in Female Cells. J Bone Miner Res, 2021. 36(6): p. 1104–1116.

47. Chen, C. and D.N. Kalu, Strain differences in bone density and calcium metabolism between C3H/HeJ and C57BL/6J mice. Bone, 1999. 25(4): p. 413–20.

48. Ericsson, A.C. and C.L. Franklin, The gut microbiome of laboratory mice: considerations and best practices for translational research. Mamm Genome, 2021. 32(4): p. 239–250.

49. Gozalo, A.S. and W.R. Elkins, A Review of the Effects of Some Extrinsic Factors on Mice Used in Research. Comp Med, 2023. 73(6): p. 413–431.

50. Reul, O.N., et al., Practical Considerations for the Design, Execution, and Interpretation of Studies Involving Whole-Bone Bending Tests of Rodent Bones. J Vis Exp, 2023(199).

51. Nyman, J.S., et al., Predicting mouse vertebra strength with micro-computed tomography-derived finite element analysis. Bonekey Rep, 2015. 4: p. 664.

